# Mesoscale medial temporal lobe connectivity patterns relate to tau pathology and memory in older adults

**DOI:** 10.64898/2026.08.18.745399

**Authors:** Larissa Fischer, Niklas Vockert, Joseph Höpker Fernandes, Berta Garcia-Garcia, Sebastian N. Roemer-Cassiano, Nicolai Franzmeier, Helena M. Gellersen, Beate Schumann-Werner, Niklas Behrenbruch, Svenja Schwarck, Eóin N. Molloy, Gusalija Behnisch, Constanze Seidenbecher, Björn H. Schott, Barbara Morgado, Hermann Esselmann, Jens Wiltfang, Henryk Barthel, Osama Sabri, Michael C. Kreissl, Emrah Düzel, Stefanie Schreiber, Esther Kuehn, Anne Maass

## Abstract

The medial temporal lobe (MTL) is crucial for episodic memory. Tau pathology is a hallmark of Alzheimer’s disease (AD) and accumulates in layer-specific patterns in the MTL during aging. It is, however, unclear whether early AD pathology relates to mesoscale network signatures distinct from non-pathological aging. To address this gap, we acquired 7 Tesla submillimeter-resolution resting-state fMRI, plasma-based AD biomarkers, glial fibrillary acidic protein (GFAP) levels, *APOE* genotype, regional [^18^F]PI-2620 tau PET burden, and longitudinal episodic memory data in 75 cognitively unimpaired older adults. Older age was associated with lower perirhinal– hippocampal connectivity and lower network segregation, whereas higher plasma-based AD pathology was associated with higher perirhinal–hippocampal connectivity. Furthermore, temporal-lobe tau burden was related to altered connectivity patterns in tau-vulnerable MTL subfields and layers, dependent on GFAP levels. Retrosplenial tau burden was associated with higher hippocampal-retrosplenial connectivity consistent with tau spread along canonical hippocampal output pathways. Finally, higher connectivity within the hippocampus attenuated the negative association between temporal-lobe tau burden and memory performance but predicted unfavorable memory trajectories. Our findings show differential associations of age and AD pathology with mesoscale MTL-connectivity patterns. Importantly, increased hippocampal connectivity may support memory function in the short term while contributing to subsequent memory decline.

## 1. Introduction

The medial temporal lobe (MTL) is an essential component of the episodic memory network^1,2^ and is affected by aging and early tau pathology. Tau tangles and amyloid-beta (Aβ) plaques are the hallmarks of Alzheimer’s disease (AD)^3,4^. According to human *in vivo* PET studies, about 15% and 35% of cognitively unimpaired older adults at age 80 are classified as biomarker positive for tau^5^ and Aβ pathology^6^, respectively, putting them at high risk for future cognitive impairment^7^. Post-mortem evidence further suggests that low but detectable AD pathology, particularly tau, can be found in the majority of cognitively unimpaired older adults^3,8^. As diverse age-related changes occur at the cellular and synaptic level^9^, differentiating the effects of aging and early AD pathology on functional brain dynamics remains challenging but is of critical importance for prevention trials.

Laminar MTL pathways have been studied extensively in rodent, monkey, and human post mortem research. Cortical information enters the hippocampus through canonical input pathways from the superficial layers II/III of the perirhinal and parahippocampal cortex to superficial entorhinal cortex layers II/III^10^. From there, input is distributed to superficial layers of the hippocampal subfields dentate gyrus, CA1, and CA3. Information within the hippocampus is then passed on between deep pyramidal hippocampal subfield layers, such as from deep CA1 to deep subiculum^11^. A prominent hippocampal output pathway originates in deep pyramidal subiculum layers and projects to the neocortex, including the neighboring retrosplenial cortex (RSC)^12–14^. Given human post mortem evidence that early tau pathology affects superficial entorhinal cortex layer II, CA1, and subiculum^3,15^, the earliest signs of AD pathology could be detectable via specific functional disturbances in this layer-specific MTL organization.

As pathology progresses, tau spreads through connected networks (e.g., from deep hippocampal output layers to the RSC), facilitated by Aβ-related neural hyperactivity^16–19^. Human studies have linked early AD pathology in cognitively unimpaired older adults to functional changes such as MTL fMRI hyperconnectivity (i.e. higher connectivity measures relative to young adults or increases over time^20^), but these changes have not been investigated at the detailed, mesoscale level. Further, the progression of AD pathology is modulated by genetic and inflammatory factors. Among those factors, the Apolipoprotein-E4 (*APOE4*) genotype is the strongest genetic risk factor for sporadic AD^21,22^ and related to heightened neuronal excitability and Aβ accumulation^23,24^. Another key modulator is neuroinflammation: astrocytes regulate neuronal excitability, glutamate homeostasis, and synaptic function. Plasma-based glial fibrillary acidic protein (GFAP) may be interpreted as a proxy for astrocytic reactivity^25^, and increasing astrocyte reactivity may signal a shift from neuroprotective or compensatory responses in early AD to pathology-promoting states. The mechanisms and time course of this transition, however, remain poorly understood^26,27^.

Functional changes with age and AD pathology differentially impact cognition in older adults. Human fMRI studies have reported lower or decreasing resting-state functional connectivity (rsFC)^28,29^ and less efficient information processing^30,31^ as features of normal aging. Additionally, higher activity of CA3 was reported in older adults and aged rodents^32–34^. Preserved, youth-like functional network organization in aging has been associated with better memory performance^35^. However, most human studies on normal aging do not assess Aβ or tau levels, making it difficult to disentangle effects of aging and early AD pathology. For example, higher Aβ burden has been associated with higher entorhinal–hippocampal rsFC and lower mnemonic discrimination performance in cognitively unimpaired older adults^36^. Other studies suggest that higher rsFC reflects compensatory processes supporting cognitive functioning^37–39^. Higher rsFC can be interpreted as compensatory when it attenuates the negative association between pathology and cognition^40^. However, human fMRI studies have also shown that higher rsFC predicts greater tau burden in connected regions^41,42^, consistent with trans-neural tau spread reported in animal models^43,44^. Whether higher MTL rsFC preserves cognition, accelerates cognitive decline, or both remains a key unresolved question.

While animal and post-mortem studies have identified subfield- and layer-specific vulnerability of MTL circuits, it is unclear whether aging and early AD pathology exert differential effects on MTL network organization at the subfield and laminar level. It is also underexplored how such changes may relate to cognition. Furthermore, it remains unknown whether rsFC in living humans reflects the selective susceptibility of superficial entorhinal cortex layer II to tau pathology^45,46^ and the involvement of hippocampal output layers in tau propagation to neocortex^47^. Finally, it is unclear whether GFAP and *APOE4* moderate these associations, as prior findings suggest complex interactions^48,49^. Resolving these questions requires moving beyond region-level rsFC to examine the mesoscale rsFC of subfields and layers in humans *in vivo*. Submillimeter-resolution 7T fMRI has enabled detailed subfield-level investigations of the MTL^50–52^. Furthermore, laminar connectivity in younger adults has been studied using geometry-based approaches of equally-spaced layers that were binned to superficial and deep layer compartments in MTL^53–56^ and neocortex^57,58^ However, laminar fMRI studies of MTL circuits in the context of aging and early AD pathology in cognitively unimpaired older adults are lacking.

Here, we leveraged 7T submillimeter fMRI to characterize subfield- and layer-specific rsFC within the MTL and its output pathway to RSC in a cohort of cognitively unimpaired older adults. By integrating plasma-based AD biomarkers, regional tau PET burden, *APOE4* genotype, plasma-based GFAP, and longitudinal episodic memory data, we combined multimodal phenotyping with a systematic investigation of how mesoscale MTL network signatures relate to non-pathological aging and early AD pathology (see Figure 1 for a summary). First, our findings suggest that non-pathological aging and plasma-based AD pathology have distinct associations with mesoscale MTL rsFC and network efficiency. Second, local tau burden is associated with layer-specific rsFC, with these associations moderated by GFAP levels. Third, the data indicate that higher rsFC within the hippocampal head confers initial compensation against tau-related memory deficits at the long-term cost of steeper memory decline. Overall, our findings provide *in vivo* evidence that non-pathological aging and early AD pathology differentially shape MTL connectivity patterns at the mesoscale with differential implications for cognition.

**Figure 1.**
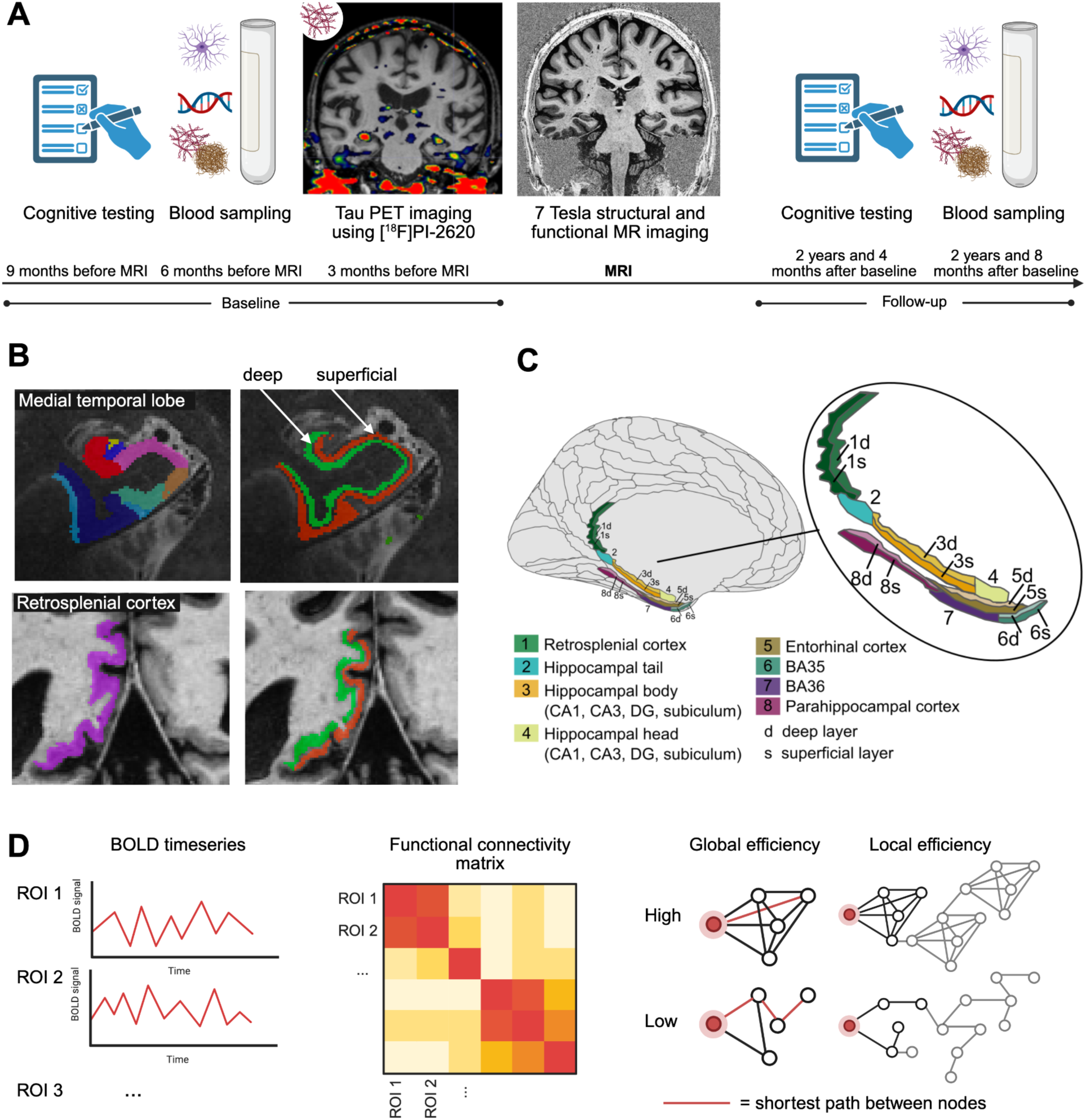
Study overview and applied methodology. **A.** During the baseline visits, cognitive testing, blood sampling, 7 Tesla submillimeter-resolution T1-, T2-, and EPI resting-state MRI, and PET imaging were performed. PET DVR images were generated from dynamic [^18^F]PI-2620 tau PET (0-60 min). An episodic memory composite built from word list recognition and word list, story, and Rey figure delayed recall was used in this study. We preregistered our study on the OSF^59^. **B.** Upper row: ASHS segmentation of the 7 Tesla T2-weighted image was used to derive medial temporal lobe (MTL) ROIs, a deep (green MTL segmentation) and a superficial layer (red MTL segmentation) was created using LayNii (for details see Supplementary Figure S1). Deep layers of CA1, CA3, and subiculum refer to the “inner” layer segment likely covering pyramidal layers and superficial layers refer to the “outer” segment likely covering stratum lacunosum moleculare. Lower row: Fastsurfer DKT atlas segmentation of the 7 Tesla T1-weighted image was used to derive a retrosplenial cortex (RSC) ROI. The same layer approach was applied. Overall, layers were derived in the hippocampal body for CA1, CA3, subiculum, as well as for BA35, entorhinal cortex, parahippocampal cortex, and RSC, where layers passed visual inspection. **C.** Investigated brain regions included MTL subregions and the RSC. For visualization purposes only, regions of interest (ROIs) from the Glasser atlas and *ggseg* were adapted in RStudio. Only the left hemisphere is shown. **D)** BOLD time-series from each ROI were extracted to form a functional connectivity matrix. ROI-to-ROI functional connectivity analysis was conducted as well as graph analysis. Three graph measures were assessed. The characteristic path length is the average shortest path length between all pairs of nodes (ROIs) of a graph (network). Global efficiency is computed as the average inverse shortest path length between all node pairs, emphasizing short paths and reflecting network integration. Local efficiency is calculated as the average efficiency of information transfer within the immediate neighborhood of a node, indexing network segregation and redundancy. All three measures were computed at the network level (averaged across all nodes) and at the nodal level (for individual nodes). Depicted are schematic examples of low and high nodal global and local efficiency for a single node (red) of a network. Grey nodes and edges are not part of the depicted node’s direct neighborhood.

## 2. Results

### 2.1. Participants

Our preregistered analyses^59^ were performed in cognitively unimpaired older adults aged 60 years and older from a deeply phenotyped longitudinal aging cohort (see Behrenbruch et al., 2025^60^). The assessment included extensive neuropsychological testing, 7T MRI, and blood sampling. As a composite score of plasma-based Aβ and tau, the AT-Term (1/ (Aβ_1-42_/Aβ_1-40_) * p- tau_217_) was used^61^. A subgroup of individuals underwent [^18^F]PI-2620 tau-PET imaging^62^ (see Figure 1A).

To address our research questions, three partially overlapping samples were defined out of this group: Sample A to investigate non-pathological aging: plasma-based Aβ- and tau negative (A-T-) participants (N=57) defined via a threshold of AT-Term ≤ 1.97^63^; Sample B to investigate plasma-based AD pathology: all participants with plasma biomarkers (N=72) defined via the AT-Term; Sample C to investigate regional tau PET burden in temporal lobe and RSC: all participants with [^18^F]PI-2620 tau PET data (N=55). Demographics are summarized in Table 1.

**Table 1:**
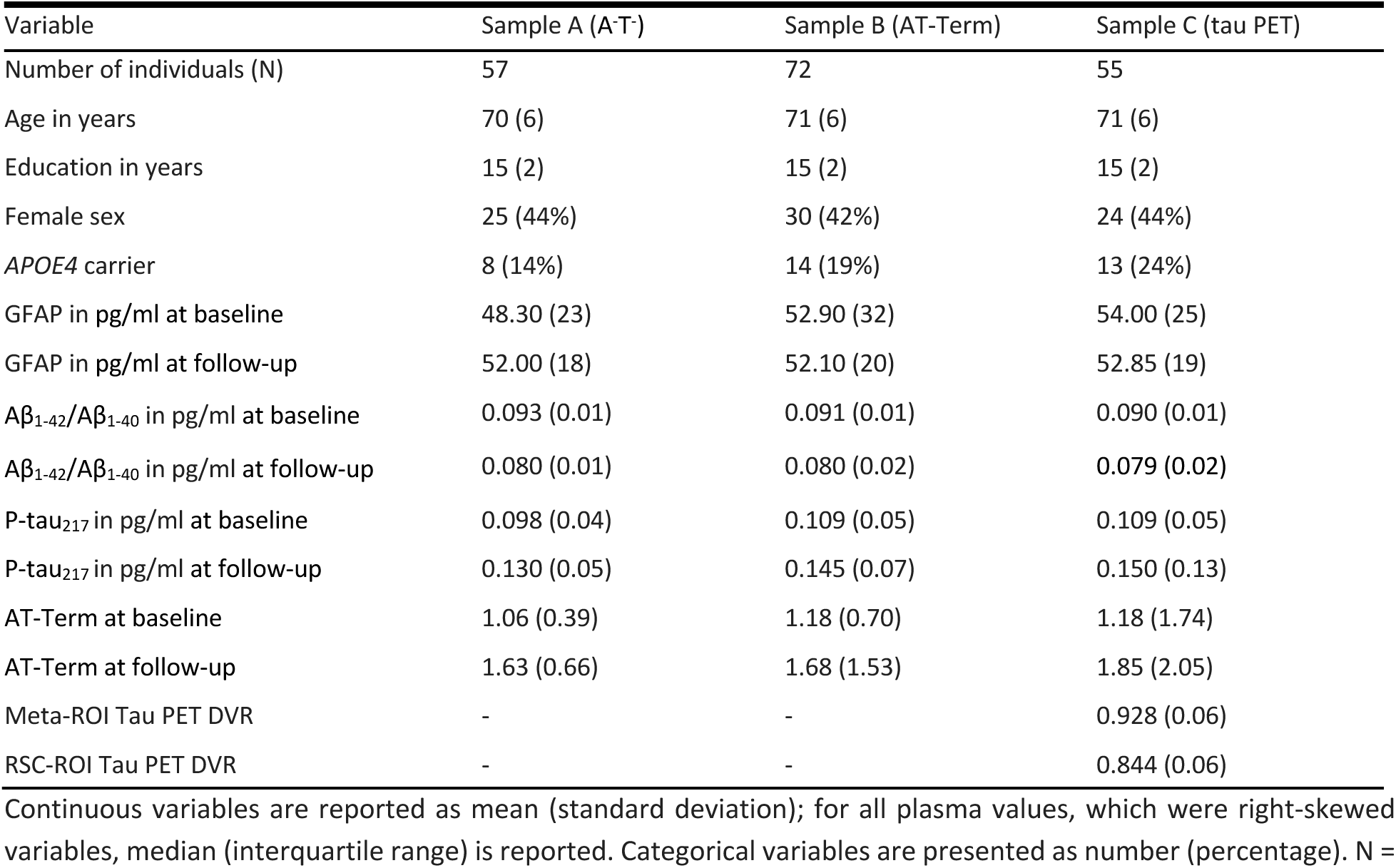
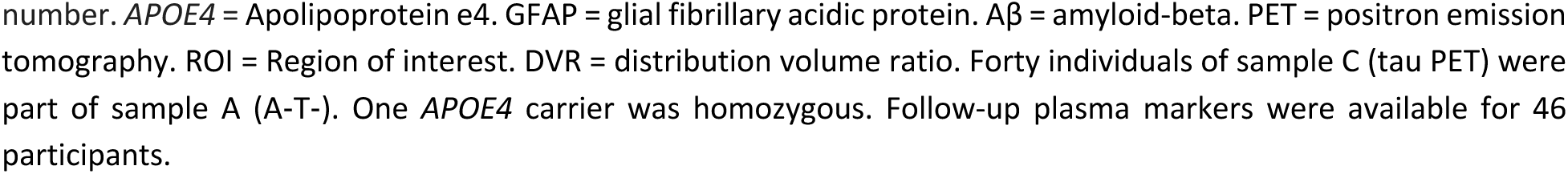
Final sample demographics after exclusions.

| Variable | Sample A (A-T-) | Sample B (AT-Term) | Sample C (tau PET) |
| --- | --- | --- | --- |
| Number of individuals (N) | 57 | 72 | 55 |
| Age in years | 70 (6) | 71 (6) | 71 (6) |
| Education in years | 15 (2) | 15 (2) | 15 (2) |
| Female sex | 25 (44%) | 30 (42%) | 24 (44%) |
| APOE4 carrier | 8 (14%) | 14 (19%) | 13 (24%) |
| GFAP in pg/ml at baseline | 48.30 (23) | 52.90 (32) | 54.00 (25) |
| GFAP in pg/ml at follow-up | 52.00 (18) | 52.10 (20) | 52.85 (19) |
| A$\beta_{1-42}$/A$\beta_{1-40}$ in pg/ml at baseline | 0.093 (0.01) | 0.091 (0.01) | 0.090 (0.01) |
| A$\beta_{1-42}$/A$\beta_{1-40}$ in pg/ml at follow-up | 0.080 (0.01) | 0.080 (0.02) | 0.079 (0.02) |
| P-tau<sub>217</sub> in pg/ml at baseline | 0.098 (0.04) | 0.109 (0.05) | 0.109 (0.05) |
| P-tau<sub>217</sub> in pg/ml at follow-up | 0.130 (0.05) | 0.145 (0.07) | 0.150 (0.13) |
| AT-Term at baseline | 1.06 (0.39) | 1.18 (0.70) | 1.18 (1.74) |
| AT-Term at follow-up | 1.63 (0.66) | 1.68 (1.53) | 1.85 (2.05) |
| Meta-ROI Tau PET DVR | - | - | 0.928 (0.06) |
| RSC-ROI Tau PET DVR | - | - | 0.844 (0.06) |

Continuous variables are reported as mean (standard deviation); for all plasma values, which were right-skewed variables, median (interquartile range) is reported. Categorical variables are presented as number (percentage). N =
number. *APOE4* = Apolipoprotein e4. GFAP = glial fibrillary acidic protein. $A\beta$ = amyloid-beta. PET = positron emission tomography. ROI = Region of interest. DVR = distribution volume ratio. Forty individuals of sample C (tau PET) were part of sample A (A-T-). One *APOE4* carrier was homozygous. Follow-up plasma markers were available for 46 participants.

### 2.2. Older age is related to lower dentate gyrus connectivity and altered CA3 network properties

First, we investigated associations between age and network measures in sample A (A^-^T^-^ individuals) to assess network dynamics in non-pathological aging. We assessed rsFC using fMRI data with a 0.9 mm isotropic voxel resolution. Given that MTL regions are approximately 2–3 mm thick in older adults^64,65^, this spatial resolution enabled the investigation of cortical depth-dependent connectivity *in vivo*. The respective BOLD time-series were extracted to conduct ROI-to-ROI rsFC analysis and the network-based statistic (NBS) for correction of multiple comparisons was applied (see Figure 1C and 1D). Higher baseline age was associated with lower rsFC in a cluster (*p*-FDR = 0.044 [95%CI*_p_* 0.040, 0.048], Figure 2A) consisting of the connections left BA36 - left dentate gyrus body (β = -0.46 [95%CI -0.71, -0.20], t(52) = -3.63, Figure 2B, Table S1) and left BA36 - right deep subiculum body (β = -0.46 [95%CI -0.72, -0.21], t(52) = -3.61, Table S2). Further, the graph metrics global efficiency, local efficiency, and characteristic path length were assessed, including Benjamini-Hochberg FDR-correction for node-level metrics. Local efficiency is a measure assessing subnetwork segregation within the immediate neighborhood of a node, either averaged across the whole network or computed for an individual node. Older age was associated with lower local efficiency of the whole network (β = -0.24 [95%CI -0.52, -0.03], t(52) = -1.75, *p* = 0.043, Table S3; Figure 2C and 2D) and lower nodal local efficiency of the left CA3 head (β = -0.66 [95%CI -0.93, -0.39], t(52) = -4.94, *p*-FDR < 0.001, Table S4).

**Figure 2.**
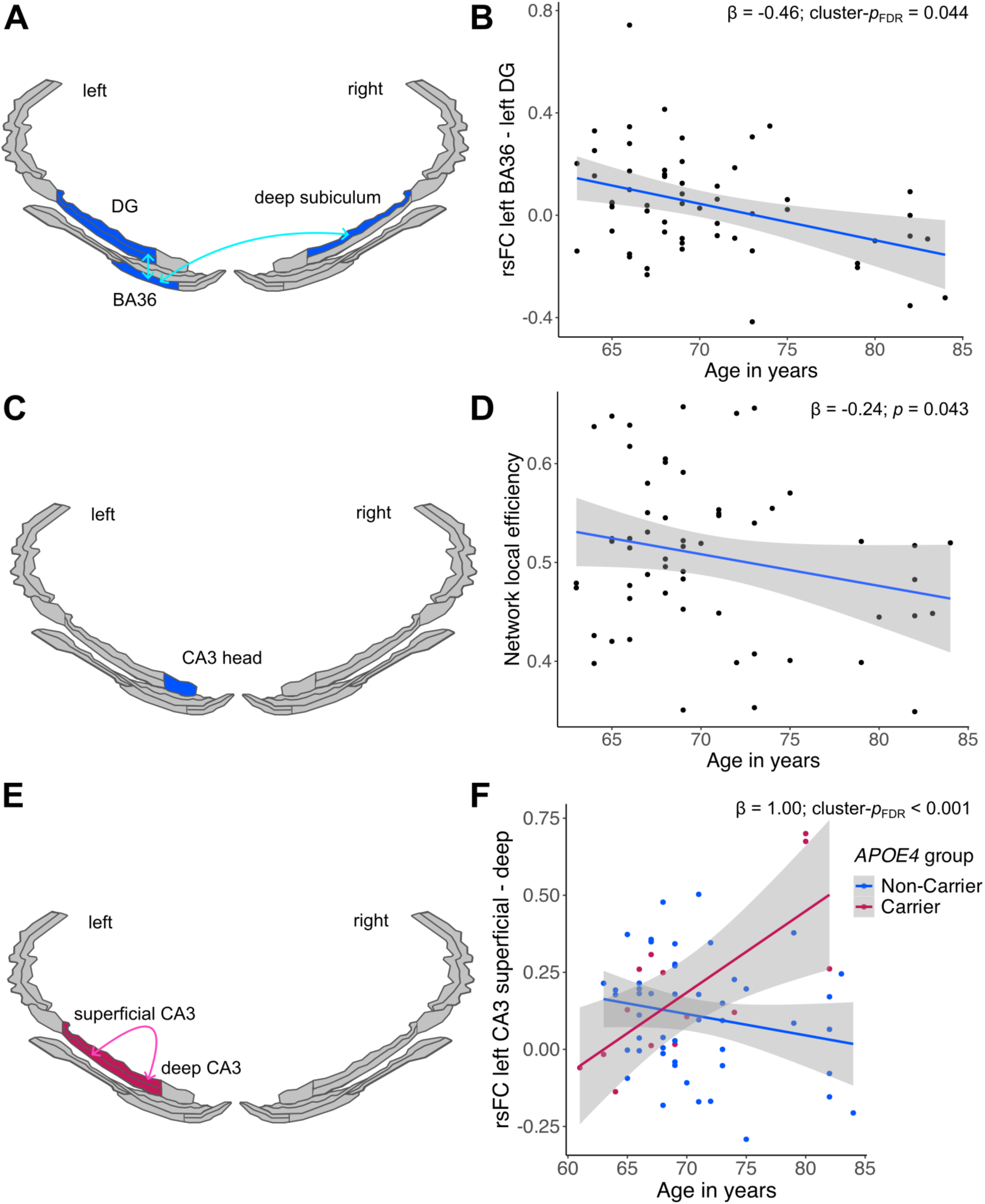
Older age is related to lower dentate gyrus connectivity and altered CA3 network properties. We investigated sample A (A^-^T^-^ individuals). For visualization purposes only, regions of interest (ROIs) from the Glasser atlas and *ggseg* were rendered in RStudio. ROIs were divided into superficial and deep layers. **A.** Higher baseline age was associated with lower resting-state functional connectivity (rsFC) between left BA36 and left dentate gyrus (DG) and between left BA36 and right deep subiculum body (highlighted in blue). **B.** The functional connection exhibiting the strongest effect (left BA36 - left DG) is used for visualization of the association of older age and lower rsFC. **C.** Higher baseline age was associated with lower local efficiency of the whole network and left CA3 head (highlighted in blue). **D.** Lower network local efficiency is associated with older age. **E.** Baseline age interacted with *APOE4* group regarding rsFC between left superficial and deep CA3 (highlighted in red). **F.** The respective functional connection (left superficial - deep CA3) is used for visualization of the interaction of age by *APOE4* group on rsFC. Older baseline age was associated with higher rsFC between left superficial and deep CA3 in *APOE4* carriers (red line), but not non-carriers (blue line).

Moreover, there was an interaction effect between age and *APOE4* group for a cluster (*p*- FDR < 0.001 [95%CI*_p_* 0.000, 0.002], Figure 2E) in the left hippocampal body, consisting of the connection superficial CA3 - deep CA3 (β = 1.36 [95%CI 0.57, 2.15], t(51) = 3.45, Cohen’s f^2^ = 0.26, Table S5). Post-hoc analyses suggest a positive age-related slope of rsFC in *APOE4* carriers (small subgroup of 8 individuals) (β = 0.52 [95%CI -0.25, 1.30], t(4) = 1.87, *p* = 0.135, Table S6), and a negative slope in *APOE4* non-carriers (β = -0.18 [95%CI -0.47, 0.11], t(45) = -1.22, *p* = 0.229, Table S7). As a sensitivity analysis, we repeated the model including all 14 *APOE4* carriers of the cohort while additionally controlling for AD pathology by including the AT-Term as a covariate. The interaction effect (*p*-FDR < 0.001 [95%CI*_p_* 0.000, 0.003] was also observed in this larger sample (β = 1.00 [95%CI 0.48, 1.52], t(56) = 3.85, Cohen’s f^2^ = 0.24, Figure 2F, Table S8) and older age was associated with higher rsFC in the group of 14 *APOE4* carriers (β = 0.71 [95%CI 0.12, 1.30], t(9) = 2.72, *p* = 0.024, Table S9). There was neither an interaction effect between age and GFAP on rsFC, nor between age and *APOE4* group or age and GFAP on graph measures (all *p* > 0.05).

### 2.3. Higher and increasing plasma-based Alzheimer’s disease pathology burden is related to higher connectivity and lower global integration

We next investigated associations between plasma-based AD biomarkers and network measures in sample B (AT-Term) to determine whether AD pathology shows distinct associations from those observed for non-pathological aging. As the AT-Term is right-skewed, analyses were conducted using log-transformed values. Overall, the baseline AT-Term was higher with older age (β = 0.13 [95%CI 0.04, 0.21], t(67) = 2.94, *p* = 0.004) and higher in *APOE4* carriers compared to non-carriers (β = 0.40 [95%CI 0.19, 0.61], t(67) = 3.78, *p* < 0.001). A higher baseline AT-Term was also associated with higher rsFC in a cluster (*p*-FDR = 0.042 [95%CI*_p_* 0.038, 0.046], Figure 3A) consisting of the connection right BA36 - left CA1 head (β = 1.13 [95%CI 0.55, 1.71], t(67) = 3.91, Figure 3B, Table S10).

**Figure 3.**
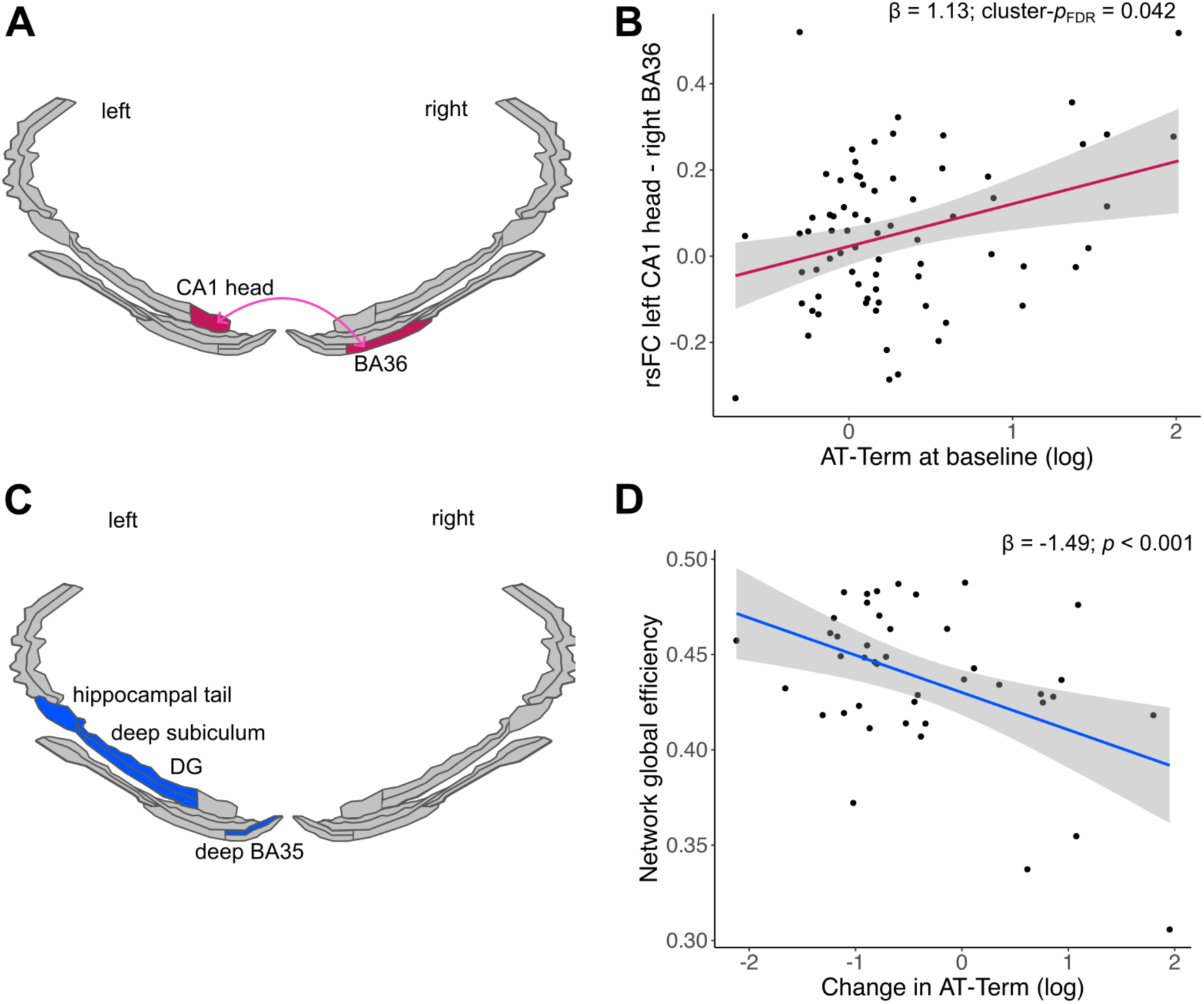
Higher and increasing plasma-based Alzheimer’s disease pathology burden is related to higher connectivity and lower global integration. We investigated sample B (AT-Term). For visualization purposes only, regions of interest (ROIs) from the Glasser atlas and *ggseg* were rendered in RStudio. ROIs were divided into superficial and deep layers. **A.** Higher baseline AT-Term was associated with higher resting-state functional connectivity (rsFC) between right BA36 and left CA1 head (highlighted in red). **B.** The respective functional connection (right BA36 - left CA1 head) is used for visualization of the association of higher AT-Term and higher rsFC. **C.** Higher change in AT-Term was associated with lower global efficiency of the network and specific nodes (highlighted in blue). All regions were located in the left hemisphere. **D.** The lower network global efficiency is used for visualization of the association of increase in AT-Term and lower global efficiency.

Longitudinal analyses showed that the AT-Term increased from baseline to follow-up, which was on average 2 years and 8 months later (days: mean=977, SD=87, mean change = 0.98 [95%CI 0.53, 1.42], t(45) = 4.42, *p* < 0.001). A larger longitudinal increase in the AT-Term was associated with higher rsFC between right BA36 - left CA1 head (β = 1.18 [95%CI 0.49, 1.86], t(41) = 3.49, Table S11), consistent with the baseline findings. However, this association did not survive cluster correction and no significant clusters were observed. Regarding graph measures, an increasing AT-Term was associated with lower global efficiency of the whole network (β = -1.49 [95%CI -2.22, -0.75], t(41) = -4.09, *p* < 0.001, Figure 3C and 3D, Table S12), lower global efficiency of hippocampal tail (which is dominated by CA1^66^), deep CA1 body, dentate gyrus body, and deep BA35 (all left hemisphere), and higher characteristic path length for right superficial BA35 (Figure 3C; see Table 2 for statistics). Global efficiency quantifies how efficiently information is exchanged between a node and all other nodes in the investigated network, reflecting its level of global integration. Characteristic path length reflects the average shortest path between a node and all other nodes, with higher or longer path lengths indicating less direct and efficient communication across the network. Both measures were computed for individual nodes and averaged across the whole network.

**Table 2.** Associations of graph measures for specific nodes and change in AT-Term.

| Graph measure | Node | beta | 95% CI | df | t | p | p-FDR |
| --- | --- | --- | --- | --- | --- | --- | --- |
| **Nodal global efficiency** | Left hippocampal tail | -0.45 | -0.72, -0.17 | 41 | -3.32 | 0.002 | 0.033 |
|  | Left dentate gyrus body | -0.48 | -0.78, -0.17 | 41 | -3.14 | 0.003 | 0.033 |
|  | Left CA1 deep body | -0.46 | -0.77, -0.15 | 41 | -2.96 | 0.005 | 0.036 |
|  | Left deep BA35 | -0.44 | -0.75, -0.13 | 41 | -2.82 | 0.007 | 0.038 |
| **Nodal characteristic path length** | Right superficial BA35 | 0.60 | 0.29, 0.91 | 41 | 3.94 | 0.001 | 0.015 |

Standardized betas, degrees of freedom (df), t-values (t), the corresponding 95% confidence interval (CI), and the FDR-corrected p-value for each node for the respective graph measure are reported. AT-Term values were log-transformed.

Neither *APOE4* group nor GFAP levels moderated the associations of baseline AT-Term or change in AT-Term with rsFC or graph-theoretical measures(all *p* > 0.05).

### 2.4. Higher tau PET burden is related to higher connectivity of susceptible MTL regions and layers

We then investigated associations between regional tau PET burden and network measures in sample C (tau PET) to identify subfield- and layer-specific connectivity signatures associated with tau pathology. [^18^F]PI-2620 distribution volume ratio (DVR) in the temporal lobe (meta-ROI) and RSC were not significantly associated with age or *APOE4* group (all *p* > 0.05), but a higher plasma-based AT-Term was associated with higher tau PET burden in the temporal lobe (β = 0.19 [95%CI 0.06, 0.30], t(49) = 2.35, *p* = 0.023) and in RSC (β = 0.16 [95%CI 0.04, 0.26], t(49) = 2.01, *p* = 0.042).

We first investigated associations between temporal lobe tau PET DVR and rsFC. There was no main effect of temporal-lobe tau burden, but there was an interaction effect between temporal-lobe tau burden and GFAP level for a cluster (*p*-FDR = 0.013 [95%CI*_p_* 0.011, 0.015], Figure 4A) including superficial entorhinal cortex, CA1, and subiculum of the left hemisphere (see Table 3 for statistics). The strongest association was observed for left CA1 head - left subiculum head (Figure 4B, Table S13). The direction of the tau–rsFC association varied with GFAP level, with the slope being positive at higher and negative at lower GFAP values when using a post-hoc median split. Regarding graph measures, higher temporal-lobe tau burden was associated with lower nodal local efficiency of right superficial entorhinal cortex, which, however, does not survive FDR correction (β = -0.45 [95%CI -0.74, -0.15], t(40) = -3.03, *p*-FDR = 0.091, Table S14).

**Figure 4.**
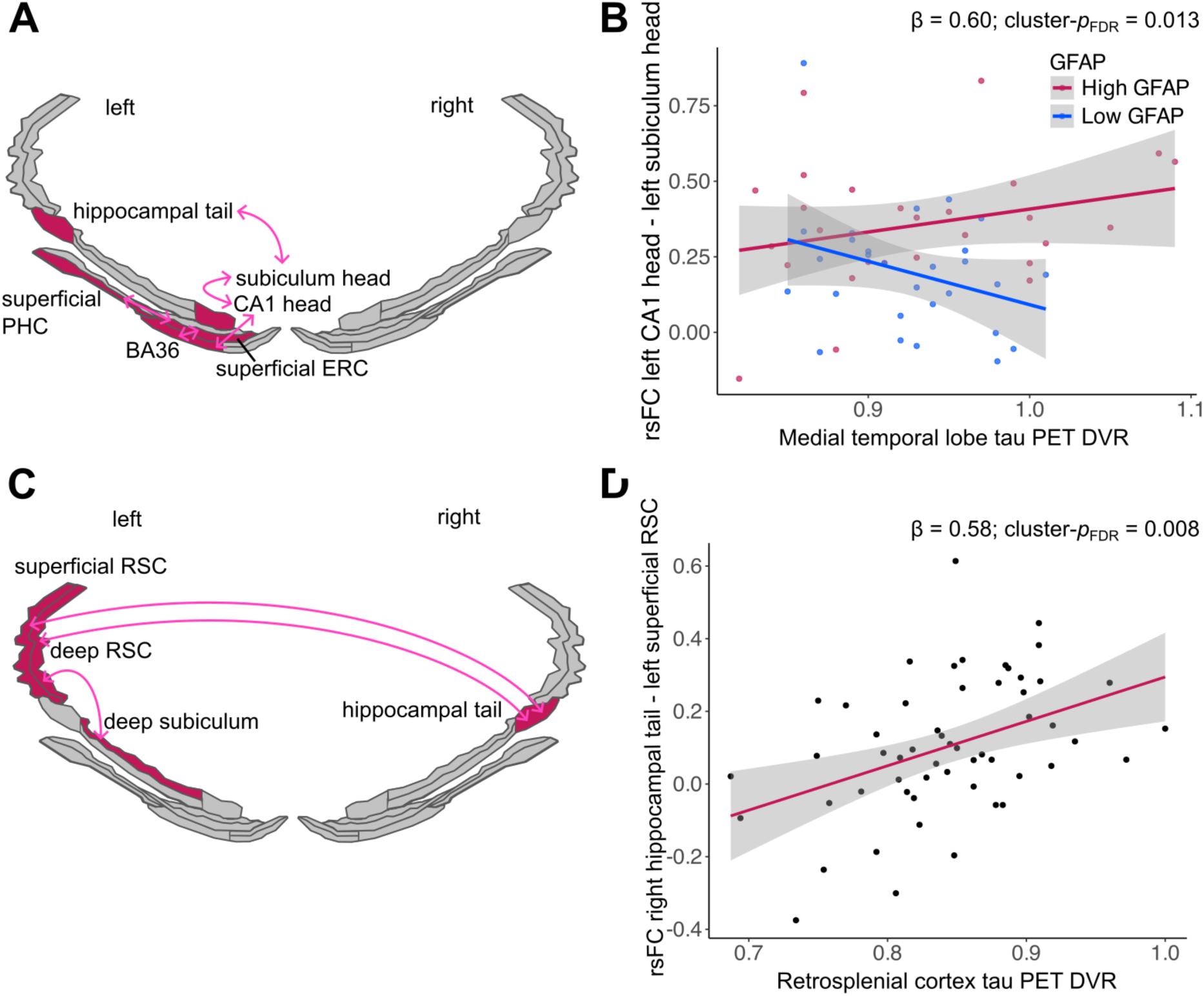
Higher tau PET burden is related to higher connectivity of susceptible MTL regions and layers. We investigated sample C (tau PET). For visualization purposes only, regions of interest (ROIs) from the Glasser atlas and *ggseg* were rendered in RStudio. ROIs were divided into superficial and deep layers. **A.** Higher temporal-lobe tau burden (i.e. [^18^F]PI-2620 DVR) was associated with higher rsFC between specific MTL regions in the presence of higher GFAP levels, highlighted in red. These regions encompass BA36, superficial PHC, superficial entorhinal cortex, CA1 head and tail, and subiculum head. All regions were located in the left hemisphere. **B.** A significant temporal-lobe tau by GFAP interaction on rsFC was observed, such that the association between temporal-lobe tau and rsFC changed from negative to positive as GFAP levels increased. For visualization, a median split for GFAP and the connection with the strongest association was used. **C.** Higher RSC tau PET burden was associated with higher rsFC between hippocampal tail and subiculum with RSC, highlighted in red. **D.** For visualization, the connection with the strongest association (right hippocampal tail - left superficial RSC) was used for visualization of the association of higher RSC tau burden and higher rsFC.

**Table 3.** Associations of functional connectivity strength and tau pathology burden.

| Model | Functional connection | beta | 95% CI | df | t |
| --- | --- | --- | --- | --- | --- |
| **Temporal-lobe tau \***<br>**GFAP** | left CA1 head - left subiculum head | 0.60 | 0.31, 0.89 | 48 | 4.13 |
|  | left BA36 - left CA1 head | 0.59 | 0.29, 0.90 | 48 | 3.89 |
|  | left superficial PHC - left superficial ERC | 0.56 | 0.24, 0.88 | 48 | 3.49 |
|  | Left hippocampal tail - left subiculum head | 0.52 | 0.2, 0.84 | 48 | 3.27 |
|  | left BA36 - left superficial ERC | 0.52 | 0.20, 0.84 | 48 | 3.26 |
| **RSC tau** |  |  |  |  |  |
|  | right hippocampal tail - left superficial RSC | 0.58 | 0.31, 0.85 | 50 | 4.26 |
|  | left deep subiculum body - left deep RSC | 0.54 | 0.26, 0.82 | 50 | 3.89 |
|  | right hippocampal tail - left deep RSC | 0.47 | 0.21, 0.73 | 50 | 3.60 |

For all functional connections that were part of significant clusters using the network-based statistics (NBS) approach, standardized betas, degrees of freedom (df), t-values (t), and the corresponding 95% confidence interval (CI) are reported. ERC = entorhinal cortex. RSC = retrosplenial cortex. PHC = parahippocampal cortex.

There was no interaction effect between temporal-lobe tau and *APOE4* group regarding rsFC, and there was no interaction effect between temporal-lobe tau burden and *APOE4* group or between temporal-lobe tau burden and GFAP regarding graph measures (all *p* > 0.05).

Second, we investigated associations between RSC tau PET DVR and rsFC. Higher RSC tau burden is associated with higher rsFC in a cluster (*p*-FDR = 0.008 [95%CI*_p_* 0.006, 0.01], Figure 4C) involving hippocampal tail (dominated by CA1^66^), deep subiculum body, and RSC, with the strongest association for rsFC between right hippocampal tail - left superficial RSC (Figure 4D, Table S15), followed by left deep subiculum body - left deep RSC, and right hippocampal tail - left deep RSC (see Table 3 for statistics).

Regarding graph measures, higher RSC tau PET burden was associated with higher nodal global efficiency (β = 0.50 [95%CI 0.24, 0.75], t(50) = 3.84, *p*-FDR = 0.014, Table S16) and shorter characteristic path length (β = -0.60 [95%CI -0.86, -0.34], t(50) = -4.67, *p*-FDR < 0.001, Table S17) for left superficial RSC.

There was no interaction effect of RSC tau by *APOE* or GFAP regarding rsFC and there was no interaction effect of RSC tau by *APOE4* group or by GFAP regarding graph measures (all *p* > 0.05).

### 2.5. Hippocampal connectivity patterns are consistent with compensation against tau-related memory deficits and subsequent memory decline

We then investigated associations between episodic memory performance and network measures to assess potential compensatory effects of altered functional dynamics in the face of pathology. We used an episodic memory composite score to investigate how rsFC and pathology measures relate to memory performance at baseline and its change over time. To assess relationships with rsFC, we limited our analyses, as defined in our preregistration^59^, to the connections that showed the strongest effect in the models described above. We hypothesized that lower rsFC would be associated with lower or decreasing episodic memory performance in non-pathological aging. Furthermore, if higher rsFC reflects resilience-related/ compensatory processes, rsFC would be expected to moderate the relationship between pathology burden and memory performance, such that higher rsFC attenuates the negative association between pathology and memory^67^.

In sample A (A^-^T^-^ individuals), baseline episodic memory performance was not significantly associated with rsFC between left BA36 - left dentate gyrus body (*p* > 0.05). However, better episodic memory was associated with higher rsFC between left CA3 superficial body - left CA3 deep body (β = 0.32 [95%CI 0.09, 0.56], t(50) = 2.75, *p* = 0.008, Figure 5A, Table S18). Better episodic memory was further associated with higher education (β = 0.42 [95%CI 0.17, 0.67], t(50) = 3.37, *p* = 0.001, Table S18) and younger age (β = -0.47 [95%CI -0.70, -0.23], t(50) = -3.95, *p* < 0.001, Table S18), as expected.

**Figure 5.**
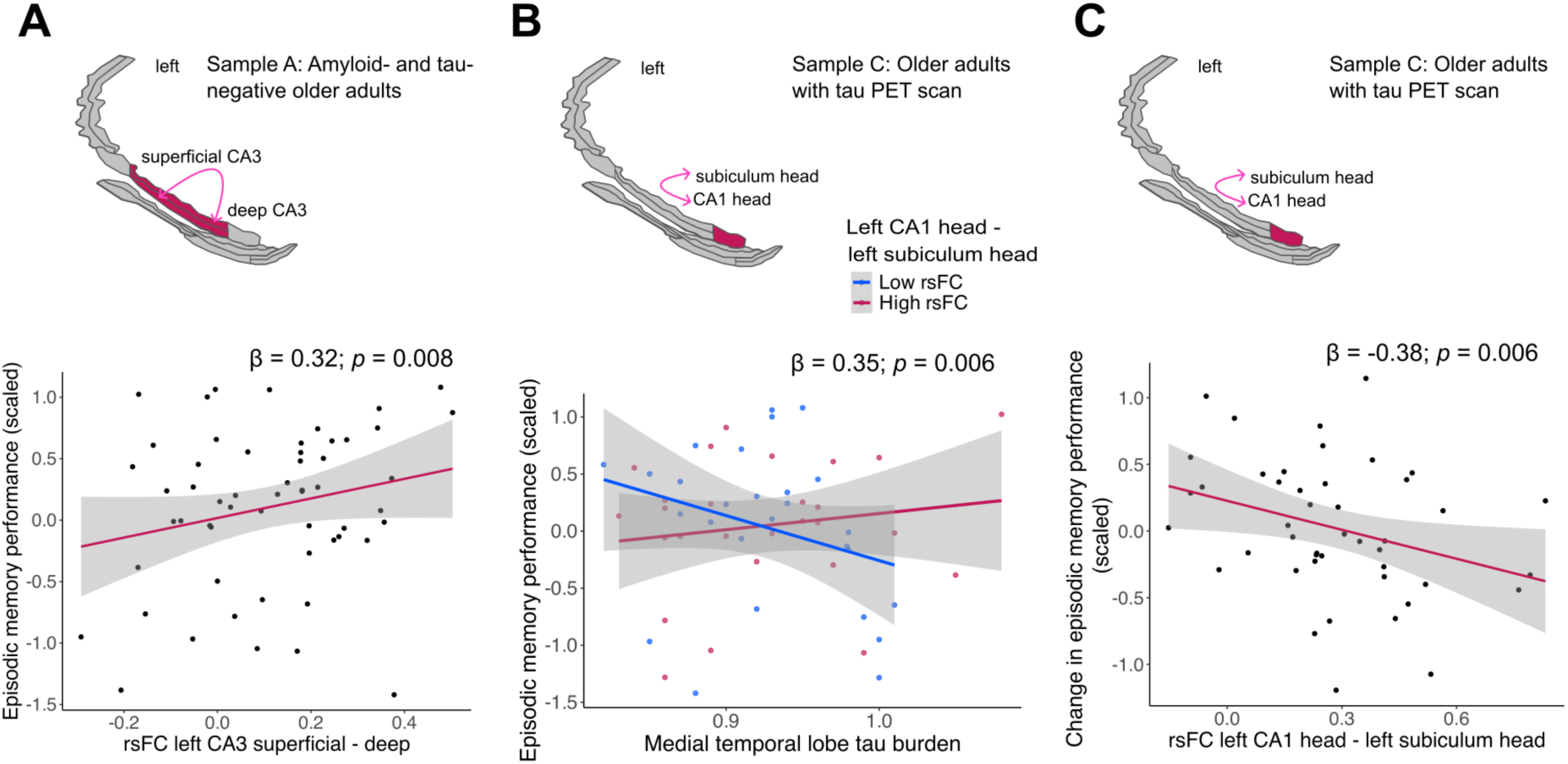
Associations between connectivity, pathology and episodic memory. For visualization purposes only, regions of interest (ROIs) from the Glasser atlas and *ggseg* were rendered in RStudio. ROIs were divided into superficial and deep layers. As preregistered ^59^, to avoid multiple testing, we assessed the relationship of episodic memory performance (baseline and change) and the connection showing the strongest association in the respective model regarding age or pathology. **A.** Sample A (A^-^T^-^ individuals). Upper panel: Older age was associated with higher resting-state functional connectivity (rsFC) between left superficial and deep CA3 in *APOE4* carriers, but not non-carriers, highlighted in red (from Figure 2E). Lower panel: Higher rsFc strength of this connection was related to better episodic memory performance. **B.** Sample C (tau PET). Upper panel: Higher temporal-lobe tau burden was associated with higher rsFC between left CA1 head and left subiculum head under high GFAP levels (strongest association), highlighted in red (from Figure 4A/B). Lower panel: The negative association between temporal-lobe tau burden and episodic memory was moderated by rsFC between left CA1 head and left subiculum head, with a weaker (less negative) association at higher rsFC (red line) compared to lower rsFC (blue line). For visualization, a median split for rsFC strength was used. **C.** Sample C (tau PET). Upper panel: Higher temporal-lobe tau burden was associated with higher rsFC between left CA1 head and left subiculum head under high GFAP levels (strongest association), highlighted in red. Lower panel: This connection shows an effect of rsFC on change in episodic memory performance, with higher rsFC being related to a steeper decline in episodic memory performance irrespective of baseline tau burden.

In sample B (AT-Term) and C (tau PET), we first assessed whether AD pathology was associated with episodic memory performance. If this was the case, we included the respective rsFC measure as moderator (interaction of pathology by rsFC), consistent with the resilience and reserve framework^40^. If no memory-pathology association was present, we included the respective rsFC measure as an additional predictor in the model to test whether it explained variance^40^. As described in the sections above, interactions with *APOE4* group and GFAP were also tested.

In sample B (AT-Term), baseline episodic memory performance was not associated with the AT-Term or, when included in the model as a predictor, with rsFC between right BA36 - left CA1 head (all *p* > 0.05). In sample C (tau PET), lower episodic memory performance at baseline was associated with higher temporal-lobe tau burden (β = -0.28 [95%CI -0.55, -0.02], t(49) = - 2.16, *p* = 0.036, Table S19). Including the interaction of rsFC between left CA1 head - left subiculum head by temporal-lobe tau burden in the model revealed that the negative pathology- memory relationship was attenuated at higher hippocampal rsFC (β = 0.35 [95%CI 0.11, 0.60], t(47) = 2.88, *p* = 0.006, Cohen’s f^2^ = 0.16, Figure 5B, Table S20). Follow-up analyses using a median-split for rsFC showed that greater temporal-lobe tau burden was associated with lower episodic memory performance in individuals with lower rsFC (β = -0.45 [95%CI -0.89, 0.00], t(22) = -2.09, *p* = 0.048, Table S21), whereas no such association was observed in individuals with higher rsFC (β = 0.13 [95%CI -0.29, 0.55], t(22) = 0.65, *p* = 0.520, Table S22). Episodic memory performance was not associated with RSC tau burden. Likewise, when included as an additional predictor in the model, rsFC between right hippocampal tail - left superficial RSC was not associated with episodic memory performance (all *p* > 0.05). There were also no significant interactions of rsFC by *APOE4* group or rsFC by GFAP levels for episodic memory performance.

Across all 61 participants with cognitive follow-up data, the episodic memory composite did not change significantly from baseline to follow-up, which was conducted on average 2 years and 4 months later (days: mean=833, SD=109) (mean change = 0.03 [95%CI -0.11, 0.17], t(60) = 0.40, *p* = 0.693). In models conducted analogously to those described for baseline performance, neither rsFC as an additional predictor nor its interactions with age, the AT-term, or RSC tau burden were significantly associated with change in performance (all *p* > 0.05). However, in sample C (tau PET; 45 individuals with cognitive follow-up data), higher baseline rsFC between left CA1 head - left subiculum head was associated with a steeper decline in episodic memory performance independent of baseline temporal-lobe tau burden (β = -0.38 [95%CI -0.65, -0.11], t(38) = -2.88, *p* = 0.006, Figure 5C, Table S23).

### 2.6. Control analyses for confounding variables, further cognitive domains, and within-sample replication

To assess further potential influencing factors, we first included white-matter DVR (off-target tracer binding) and hippocampal volume (as proxy of early atrophy) in the reported models, which did not alter the results. Additionally, hippocampal volume tended to be smaller with older age, which did not reach significance (β = -0.33 [95%CI -0.71, 0.06], t(47) = -1.72, *p* = 0.09, Table S24), but was not associated with pathology or any other measure (all p > 0.05).

Second, to assess the specificity of associations of episodic memory, pathology, and rsFC, we investigated two additional cognitive measures (see section 4.3). There were no associations of the Symbol Digit Modalities Test (SDMT) (attention and processing speed) and Cognitive Age Gap (CAG) (global cognition) with rsFC, for any of the reported models regarding cognition (all *p* > 0.05).

Third, to assess whether the resting-state results could be replicated within our sample, we used breathing-task fMRI data (with task effects regressed out), which was acquired during the same scanning session (see “Methods” section for details). When investigating the connections where associations were observed for the resting-state data, all models revealed the same directionality of effects, showing an overall similar pattern of associations, in the breathing-task fMRI data. However, effects were smaller and not all models reached significance (see Table S25 for statistics). Further, the same analyses as described above for resting-state data were conducted with the breathing-task data, including cluster correction. Models involving change in AT-Term and temporal-lobe tau PET reached significance, with connections involving CA1, subiculum, and dentate gyrus showing higher rsFC with increasing AT-Term, and connections involving subiculum and CA1 showing higher rsFC with higher temporal-lobe tau burden and GFAP levels (see Table S26 for statistics).

## Discussion

In this multimodal study, we investigated subregion- and layer-specific connectivity patterns within MTL circuits in cognitively unimpaired older adults. Three principal findings emerged. First, non-pathological aging and plasma-based AD pathology showed distinct associations with MTL network organization. For perirhinal–hippocampal pathways, older age was associated with lower rsFC and lower network segregation, whereas higher plasma-based AD pathology was associated with higher rsFC. Second, MTL [^18^F]PI-2620 tau PET burden was related to rsFC in subfields and layers expected to be vulnerable to early tau pathology. Furthermore, rsFC of hippocampal output regions and layers with RSC was related to RSC tau burden, consistent with trans-neuronal tau spread along canonical pathways. Third, higher rsFC within the hippocampal head showed a pattern suggestive of a compensation–deterioration trade-off: it attenuated the negative association of tau burden and episodic memory at baseline, but predicted memory decline over approximately two years.

In plasma-based Aβ- and tau-negative adults, we observed age-related network alterations reflecting local network dedifferentiation. Older age was associated with lower rsFC between BA36 and the hippocampal body (dentate gyrus and deep subiculum). This finding likely reflects reduced functional coupling within the broader perirhinal–hippocampal network^68–70^, as the anatomical pathways providing input to the hippocampus and feedback to perirhinal cortex both pass through the entorhinal cortex^71–73^. Our findings extend prior work on age-related lower and decreasing perirhinal–hippocampal connectivity^29,74–76^ and are consistent with the selective vulnerability of the dentate gyrus to non-pathological aging^77–79^. Furthermore, older age was associated with lower local efficiency across the network, particularly for CA3 head. Reduced local efficiency indicates lower subnetwork segregation and diminished fault tolerance^80^: If a vulnerable node becomes dysfunctional, fewer redundant connections are available, thereby exacerbating the impact of localized disruption. Additionally, older age was associated with higher within-CA3 rsFC in *APOE4* carriers but not in non-carriers. Caution should be taken here due to the small number of *APOE4* carriers. Nonetheless, our finding is consistent with prior reports of higher CA3 activity with aging, particularly in *APOE4* carriers^24,32,81,82^. Mechanistically, early age-related dentate gyrus dysfunction may reduce inhibitory control over CA3, contributing to increased CA3 self-excitation, amplifying the recurrent autoassociative circuitry of CA3, despite network-level dedifferentiation^83–85^.

We then investigated all participants with plasma-based AD biomarkers and observed pathology-related alterations indicative of network dysregulation. A higher plasma-based composite score, reflecting mainly Aβ pathology^86,87^, was associated with higher rsFC between BA36 and CA1 head. Consistent with our result, early Aβ pathology has been related to neuronal hyperexcitability. Animal studies further suggest that this hyperexcitability is related to interneuron dysfunction and disrupted synaptic transmission, leading to excitation-inhibition- imbalance^88–90^. Furthermore, an increasing AT-Term was associated with lower global network efficiency, particularly in regions known for early tau accumulation including BA35 and CA1^15^, indicating degraded integration within the MTL network. This may reflect network-level consequences of emerging synaptic dysfunction, as reported in recent studies: Aβ and tau have been associated with deficits in glutamatergic NMDA receptor function, resulting in impairments in cortico-hippocampal circuits in mice^91^. In addition, Aβ-related hyperexcitability in CA1 pyramidal neurons has been linked to altered dendritic architecture^92^. Overall, we demonstrate that non-pathological aging and early AD are associated with distinct alterations in MTL connectivity patterns at the subfield level, extending prior region-level findings^28,29^.

In participants with [^18^F]PI-2620 tau PET data, temporal-lobe tau burden was related to layer-specific rsFC in a pattern that closely mirrors the known vulnerability and propagation sequence of tau pathology reported in histological studies^93–95^. Animal models show that early local tau burden alters MTL networks by inducing neuronal hypoactivity but also by enhancing neuronal synchrony^96–98^. In our study, higher temporal-lobe tau burden was associated with higher rsFC, but only at relatively higher GFAP levels within our cohort. Specifically, superficial entorhinal cortex showed altered rsFC with superficial parahippocampal cortex and BA36. This could reflect enhanced coupling of the entorhinal cortex with its cortical input regions^10,99^ in the presence of higher tau burden and early astrocytic reactivity^97^.

Further, we observed higher rsFC between CA1 and subiculum head, two hippocampal subfields particularly vulnerable to tau accumulation^15^. Notably, we did not observe altered rsFC for the monosynaptic perforant path connection between superficial entorhinal cortex and CA1, despite this connection being particularly vulnerable to early AD-related pathology^100^. Nevertheless, we found higher rsFC for the indirect connection between BA36 and CA1, which is mediated via the entorhinal cortex^71,72^. The GFAP moderation of the tau-rsFC associations is noteworthy, as neuroinflammation, particularly astrocytic reactivity, has been recognized as a modulator of AD pathophysiology^101^. Astrocytes are regulators of neuronal function, yet the transition from a supportive role to contributing to pathology remains poorly understood, particularly in the early stages of AD^26,27^. Higher GFAP levels might reflect an initial compensatory mechanism in the face of tau pathology, promoting neuronal energy supply, or it might reflect pathological processes, potentially driving neuronal hyperactivation^26,102–104^. Whether early GFAP is indicative of a neuroprotective or pathology-promoting astrocyte state remains unresolved^105–108^, though the cognitive implications discussed below suggest an involvement in initially protective mechanisms in cognitively unimpaired individuals.

When extending our analyses from temporal to neocortical tau burden, greater RSC tau burden was associated with higher rsFC of hippocampal output pathways and RSC^12,14,109^. RSC represents one of the earliest neocortical targets of hippocampal tau spread, where deep-layer pyramidal neurons are preferentially affected^47,110^. Studies in humans have demonstrated that tau accumulates preferentially along functional connections, with higher rsFC between regions predicting greater tau burden^17,41,42^, and in rodent models, tau release and propagation are neuronal-activity dependent^43,44^. Our findings may hint towards tau spreading along anatomically defined hippocampal output pathways, consistent with laminar directional connectivity^111^: Deep but not superficial CA1 and subiculum layers project prominently to the RSC^12–14^. Additionally, graph measures suggest that superficial RSC is more tightly integrated into the MTL network, rather than disconnected, in the presence of early RSC tau burden. Consistently, a previous study reported a tau-related disconnection between hippocampus and anterior cortical networks, but not posteromedial cortical networks in cognitively unimpaired older adults^112^. Whether our findings reflect compensatory network activity in response to early tau, a network-based facilitation of tau spread, or both remains an open question^113^ that will require longitudinal MRI and PET data to address.

The cognitive implications of these observed network alterations point to a differentiation between non-pathological aging and early tau pathology as well as between cross-sectional and longitudinal outcomes. In A^-^T^-^ older adults, higher within-CA3 rsFC was associated with better episodic memory. While prior reports link higher CA3 rsFC or activity to cognitive deficits in the context of early AD pathology^32,36^, our findings indicate that higher functional coupling within CA3 supports memory in non-pathological aging. In older adults with available [^18^F]PI-2620 tau PET data, higher CA1–subiculum rsFC in hippocampal head attenuated the negative association between temporal-lobe tau burden and baseline episodic memory, suggesting an initial compensatory benefit. However, higher CA1–subiculum rsFC in the hippocampal head predicted a steeper memory decline over approximately two years, irrespective of tau burden at baseline. This finding is consistent with the hypothesis that higher connectivity, while initially supporting memory, may accelerate pathology spread^43,44^ and related cognitive decline^114^. Furthermore, this finding is consistent with the broader concept that compensatory network configurations shift from adaptive to maladaptive as pathology progresses^20,115^.

Several limitations warrant consideration. RsFC is correlational and does not permit causal inference, thus, future work could employ dynamic causal modelling or similar directional modeling of functional connectivity data^116–118^. Additionally, the resting-state acquisition was limited to approximately five minutes, constraining reliability^119,120^. Nonetheless, the within-sample replication using breathing-task fMRI data showed consistent directionality of effects across models, although not all results survived cluster correction. This may reflect reduced signal-to-noise ratio due to increased motion during paced breathing, residual task-related variance despite regressing out task effects, or the additional variability introduced by the data-driven cluster-forming threshold in network-based statistics. Furthermore, our BOLD-based laminar fMRI carries interpretive constraints: Although acquired at submillimeter resolution, the effective spatial resolution of fMRI remains limited, leading to partial-volume effects and signal mixing across layer compartments^121^. Deep and superficial compartments reflect geometrically defined cortical depths rather than histologically defined anatomical layers^122^. Further, a vascular bias has been reported in deep CA1 and subiculum^54^ and vascular properties change with aging^123^. However, converging evidence from multimodal imaging and intracranial recordings support that rsFC captures meaningful neuronal dynamics^124–126^. Sequences with higher spatial specificity such as VASO should be considered for future laminar MTL work alongside NORDIC-PCA to boost SNR^55,127^.

Overall, our findings suggest that 7T laminar fMRI can resolve human MTL network alterations at a mesoscale level that are related to tau vulnerability, previously characterized through animal and post-mortem studies. By linking subfield- and layer-specific functional connectivity to plasma-based AD biomarkers, tau PET, and longitudinal episodic memory data, we provide *in vivo* human evidence for specific MTL connectivity patterns with age and early AD pathology. Longitudinal studies combining ultra high-field fMRI with PET will be essential to determine whether laminar connectivity changes precede, parallel, or accelerate tau accumulation, and to assess potentially compensatory network dynamics.

## 3. Methods

### 3.1. Participants

All 75 participants of our preregistered study^59^ were part of the aging cohort of the Z03 project within the Collaborative Research Center (CRC) 1436 “Neural Resources of Cognition”. Participants additionally participated in a subproject where 7T MRI was performed within less than a year after baseline characterization (days: mean=223, SD=98). In short, participants were cognitively unimpaired adults over the age of 60 who performed within the age-, sex- and education-adjusted normative range on the CERAD-Plus Neuropsychological Assessment Battery^128^. Additionally, they had no major age-related illnesses and normal or corrected to normal vision. For details, see Behrenbruch et al., 2025^60^ and https://sfb1436.de/. Blood markers for Aβ_1-42_, Aβ_1-40_, p-tau_217_, and GFAP, as well as *APOE* genotype and data from extensive neuropsychological testing were available. After preprocessing of the MRI data, three participants with excessive motion during the resting-state scan (more than 20% of volumes flagged as motion outliers, see section 4.5) were excluded, no participants were excluded due to major signal drop-out. Of the remaining 72 participants, 55 participants underwent [^18^F]PI-2620 tau-PET imaging. Follow-up neuropsychological data were available for a subsample of N=61 participants and took place on average 2 years and 4 months after baseline (days: mean=833, SD=109). Follow-up plasma markers were available for N=46 participants, acquired on average 2 years and 8 months after baseline (days: mean=977, SD=87).

From the remaining 72 participants, we formed three samples to address our specific research questions regarding dissociable association of AD-independent non-pathological aging and AD pathology with rsFC. Sample A consisted of plasma Aβ- and tau-negative older adults (A- T-, N=57, 70±6 years, 25 female, 8 *APOE4*) to investigate the effects of baseline age. A threshold of AT-Term ≤ 1.97 was used to define biomarker negativity. This threshold was determined via Gaussian mixture modeling for the whole Z03 cohort (N = 346, 72±8 [age range: 60 to 94 years]) as utilized previously^63^. Sample B consisted of all older adults (N=72, 71±6 years, 30 female, 14 *APOE4*) because plasma biomarkers were available for the entire cohort, enabling analyses of the effects of AD pathology quantified by the AT-Term. Sample C consisted of older adults with available tau PET (N=55, 71±6 years, 24 female, 13 *APOE4*) to investigate the effects of localized tau pathology. Demographic details can be found in Table 1.

All study procedures and experimental protocols (see also German Clinical Trials Register: DRKS00032449) were approved by the local Ethics committee of the Medical Faculty, Otto-von-Guericke University Magdeburg (200/19) in accordance with the Declaration of Helsinki. Written informed consent was obtained from all participants prior to each experimental procedure and they were financially compensated for their time. An overview of the data and measurements is provided in Figure 1A.

### 3.2. *APOE* genotyping and plasma-based biomarkers

EDTA-plasma sampling was used to determine *APOE* group, Aβ_1–40_, Aβ_1–42_, p-tau_217_, and GFAP levels. Blood was taken on average 6 months before 7T MRI.

Two EDTA plasma samples were centrifuged at 2000 × g for 10 min at room temperature. The buffy coat was extracted and stored at 4°C for genotyping. Genomic DNA was isolated using the QIAamp® DNA Blood Mini Kit. APOE genotyping was performed using a polymerase chain reaction–based approach followed by restriction fragment length polymorphism analysis. *APOE4* carriers were defined as individuals with one or two *APOE4* alleles, whereas *APOE4* non-carriers were defined as individuals without an *APOE4* allele.

The remaining plasma was aliquoted and frozen at – 80°C before Aβ_1–40_, Aβ_1–42_, p-tau_217_, and GFAP were determined using Immunoreaction Cartridges on the fully automated Lumipulse G600II System, as described in Morgado et al, 2024^61^. All assays met the manufacturer’s quality control specifications, with quality control samples analyzed before and after testing each day and assay precision below 5% coefficient of variation across all assays. Internal quality control samples were included in each run to assess inter-assay variability. All measurements were performed in single determinations in accordance with the manufacturer’s instructions. The AT-Term = 1/ (Aβ_1-42_/Aβ_1-40_) * p-tau_217_ was used to investigate a continuous composite measure of Alzheimer’s disease pathology, with higher values indicating more pathology^61^.

### 3.3. Cognitive testing

All participants underwent extensive cognitive testing. For this study, an episodic memory composite score was calculated similar to the Repeatable Battery for Assessment of Neuropsychological Status (RBANS) delayed memory index score^129^. This score integrates word-list recognition as well as delayed free recall of word lists, figures, stories. Specifically, the delayed free recall and recognition (corrected hit rate) of the Verbal Learning and Memory Test (VLMT)^130^ (word list of 15 words), the delayed free recall of the Rey Complex Figure Test and Recognition Trial (RCFT)^131^ (one complex figure), and the delayed free recall of the Logical Memory II of the Wechsler Memory Scale (WMS)^132^ (two short stories) were used in this study.

As control analysis, we investigated attention and processing speed using the Symbol Digit Modalities Test (SDMT)^133^. Further, we investigated the Cognitive Age Gap (CAG) as a global measure of cognition^60^. The score reflects multiple domains, including verbal and visual memory, working memory, processing speed, attention, executive function, and verbal fluency. Cognitive age was estimated in the whole Z03 sample of older adults (N = 346, 72±8 [age range: 60 to 94 years]) using a previously established multivariate approach^134^. CAG was calculated as the difference between predicted and chronological age, with a negative score indicating that an individual’s cognitive performance was better than expected for their chronological age. Prediction was performed with Partial Least Squares (PLS) regression, using all available cognitive test scores (n = 19) as predictors and chronological age as the response. Scores were transformed to reduce skewness and z-standardized; missing values (≤3 per subject) were imputed with the mean z-score. Tenfold cross-validation was applied to obtain out-of-sample cognitive age predictions (for the held-out 10% of participants), followed by a statistical bias correction^135^ to account for systematic over- or underestimation at the age extremes. Prediction accuracy was assessed via Pearson correlation and variance explained (R²) across folds. For details, see^60^.

### 3.4. MRI data acquisition

7T MRI data were acquired at a Siemens MAGNETOM scanner in Magdeburg, Germany, using a Nova Medical 32-channel head coil. T1-weighted anatomical whole-brain MP2RAGE images (TR = 4800 ms; TE = 2.18 ms; TI1/TI2 = 900/2750 ms; 5**°**/3**°** flip angle; 352 slices; 17.14 min) were acquired with a 0.5 mm isotropic voxel resolution. T2-weighted turbo spin echo (TSE) images (TR = 8000 ms; TE = 92 ms; 60**°** flip angle; 50 slices; 7.38 min) were acquired with a 0.5*0.5*1.1 mm voxel resolution. The TSE protocol was optimized for MTL volumetry with 50 coronal slices orthogonal to the hippocampal long axis. FMRI data were obtained using an echo-planar imaging (EPI) sequence (TR = 2000 ms; TE = 20 ms; 80**°** flip angle; 58 slices) with a 0.9 mm isotropic voxel resolution. Resting-state scanning lasted 5.03 minutes and breathing-task scanning lasted 8.55 minutes. The breathing task comprised 10 rounds of 12s of paced breathing alternated with 30-60s of free breathing. The breathing frequency during paced breathing was set to 6s/breath, i.e. 3s breathe in and 3s breathe out, with the goal to introduce variations in the breathing patterns to assess cerebrovascular reactivity ^136^, which was not investigated in this study. The data was used for within-sample replication of the resting-state findings only (see section 4.11). In addition, 3T T1-weighted MPRAGE images were acquired during the PET-MRI scan (TR = 2500 ms, TE = 4.37 ms, TI = 1100 ms, 7° flip angle; 256 slices; 9.20 min), which were used for the preprocessing of the PET data (see section 4.8).

### 3.5. MRI preprocessing

Structural and functional 7T data were preprocessed using MATLAB, Statistical Parametric Mapping version 12 (SPM12)^137^, and image quality, segmentations, and coregistrations were visually assessed.

The structural T1 images (unified images (UNI-T1) of the MP2RAGE sequence) were segmented into gray matter, white matter, and cerebrospinal fluid. We utilized the Desikan– Killiany atlas to segment the 7T UNI-T1 image using FastSurfer^138,139^ and the 3T T1 image using FreeSurfer^140^ to derive the RSC segmentation (atlas label “isthmus cingulate”). Automated segmentation of hippocampal subfields (ASHS) was applied to the T2 image using the ASHS 7T Atlas for T2-weighted MRI^141–143^. This atlas facilitates the segmentation of hippocampal subfields and parahippocampal regions, aligning with the protocol by Berron and colleagues^141^. The automated segmentations underwent standardized visual quality control and correction, specifically targeting commonly identified errors as described before^144,145^: over- and under-segmentation of boundaries and incorrect inclusions of voxels, e.g., misidentification of the choroid plexus as part of the hippocampus or hippocampal cysts. Bilateral hippocampal volume was corrected for intracranial volume (ICV) by regressing it on ICV and adjusting relative to the sample mean ICV^146,147^.

The fMRI data were slice-time corrected, realigned to the first volume, and voxel-displacement maps were created using fieldmaps to correct susceptibility artifacts via unwarping using SPM. No smoothing was applied to preserve high anatomical specificity. Structural masks were coregistered to the EPI mean functional image using antsRegistrationSyNQuick and antsApplyTransforms^148^. For the breathing-task fMRI data, task effects were regressed out to calculate background-FC, which is used to approximate resting-state^149^ to probe within-cohort replicability of the findings (see supplementary Table S25). Outlier fMRI volumes due to excessive head motion were detected using ART^150^, where volumes with framewise displacement above 0.5 mm/ TR or a global intensity z-score of 5 were flagged, matching previous work^51,148^. fMRI sessions with more than 20% of volumes flagged as outliers were excluded from the analysis. During denoising of fMRI data using CONN, version 22a^151^, confounding effects were regressed out using realignment parameters and their first order derivatives, flagged outlier volumes, and signal from cerebral white matter and cerebrospinal fluid derived via anatomical CompCor^152,153^. A band-pass filter of 0.008 Hz to 0.09 Hz was applied to minimize noise from physiological and motion sources^154^.

### 3.6. Layer compartment processing

In the current study, the term “layers” is not used to denote histologically defined cortical or hippocampal laminae. Instead, it refers to two layered compartments (“deep layer” and “superficial layer”) that are expected to broadly align with known histological layer patterns along the basal-apical axis (from “inner” to “outer” border). In hippocampal subfields (CA1, CA3, and subiculum), the deep layer is expected to predominantly encompass pyramidal cell layers and adjacent strata, whereas the superficial layer is expected to predominantly encompass stratum lacunosum-moleculare^56^. Our approach followed a previously established protocol^155^. See Figure S1 for an overview of the layer compartment pipeline.

MTL and RSC segmentations were derived from ASHS and FastSurfer, respectively. A binarized mask of the MTL was created from the multi-label ASHS segmentations, where a one-voxel thick MTL border was defined on each slice. This was achieved by labeling all voxels that had at least one neighbouring voxel of value zero to ensure that the following rim definition exclusively affected surface voxels. For the whole MTL (without dentate gyrus) and RSC, a geometrical approach was used to define inner and outer borders on a slice-by-slice basis. For the MTL, inner and outer borders were operationally defined using directional labeling along anatomically motivated axes (inferior–superior or medial–lateral), capitalizing on the oblique-coronal T2 acquisition that is approximately orthogonal to hippocampal lamination. This procedure assigns opposing border compartments based on the first rim voxel encountered when scanning along the selected anatomical direction, with directionality adjusted by region and hemisphere to account for individual orientation differences. For the RSC, inner and outer borders were defined using an analogous approach based on the cortical geometry. All border definitions underwent manual quality control to ensure anatomical plausibility at the individual-subject level.

Layers were then assigned using the LayNii toolbox^122^. Twenty equidistant layers for the whole MTL and RSC were derived across the inner border (layer 1) to the outer border (layer 20) based on prior recommendations^111,156^. Layers 2 to 9 were binned to a deep layer compartment, layers 12 to 19 were binned to a superficial layer compartment. The superficial and the deep layer mask were then overlaid with the separate MTL ROI masks derived from ASHS. Dentate gyrus, BA36, hippocampal head and hippocampal tail were excluded from the layer formation because of insufficient visual quality control outcomes. No clear layers were distinguishable due to the complex folding structure in these areas and to no subfield delineation in the tail ^142^.

### 3.7. Functional connectivity analysis

Prior to first-level analysis of the resting-state data, for MTL (excluding CA2 due to its small size) and RSC ROIs, voxels with mean intensity over time lower than 2 standard deviations from the mean intensity (across all voxels) in an ROI were removed from the ROI^50,51,157^. For a visualization of the ROIs forming the investigated brain network, see Figure 1C. During first-level analysis, functional connectivity matrices were derived for each individual participant. Fisher z-transformed correlation coefficients for each ROI pair were derived via bivariate correlations and convolution of the equal weights of the scans with a canonical hemodynamic response function (hrf-weighting) was applied.

During second-level analysis, a separate General Linear Model (GLM) was estimated for each functional connection. Inferential statistics were performed at the network level for clusters of connections and were based on the hypothesis defining the expected association. Network-level inferences were based on nonparametric statistics from network-based statistic (NBS) analyses using randomization (10000 iterations) of residuals from the second-level model to obtain uncorrected network-level *p*-values. The results were thresholded with a combination of a cluster-forming connection-level threshold of *p* < 0.001 and a Benjamini-Hochberg^158^ false-discovery rate (FDR)-corrected network-level threshold of *p*-FDR < 0.05^151^.

### 3.8. Graph analysis

Graph analysis was performed to assess network properties, with nodes corresponding to ROIs and edges representing their functional connections^159^. We examined both network-level and node-level graph metrics to capture complementary aspects of network organization.

Network-level integration was quantified using characteristic (average) path length and global efficiency. Characteristic path length was defined as the average shortest path length between all pairs of nodes, reflecting the overall topological distance for information transfer across the network^160^. Global efficiency, computed as the average inverse shortest path length between all node pairs, provides a complementary measure of information transfer that is less sensitive to disconnected node pairs^161^. Network-level segregation was quantified using local efficiency, defined as the average inverse shortest path length between all pairs of nodes within the local neighborhood of each node, with the focal node removed, averaged across all nodes in the network^161^. At the nodal level, we assessed node-specific variants of these measures. Nodal characteristic path length was defined as the average shortest path length between a given node and all other nodes, reflecting the average topological distance of a node to the rest of the network. Nodal global efficiency was computed as the average inverse shortest path length from a given node to all other nodes, indexing how efficiently a node can exchange information with the network. Nodal local efficiency was calculated as the efficiency of the subgraph formed by a node’s immediate neighbors after removal of the focal node, capturing how efficiently information can be exchanged within the node’s local neighborhood. Together, these measures provide complementary indices of network integration^162^.

Following our preregistration^59^, the optimal cost threshold for each sample was determined by comparing global and local efficiency of real networks to random and lattice models, maximizing divergence *(Global efficiency(Data) – Global efficiency(Lattice) + Local efficiency(Data) – Local efficiency(Random)*; Supplementary Figure S2)^80^. This yielded thresholds of 0.13, 0.14, and 0.14 for samples A, B, and C, respectively, which were applied to the second-level rsFC matrices to generate binarized adjacency matrices. Statistical significance was determined with a second-level Benjamini-Hochberg FDR correction (*p*-FDR < 0.05).

### 3.9. PET data acquisition and preprocessing

PET scanning was performed 0–60 min post-injection simultaneously with 3T MRI on a Siemens Biograph mMR as described in detail in Maass et al., 2026^62^. Following an intravenous bolus of 183 MBq and a 40 mL saline flush, dynamic PET data were acquired in list-mode and reconstructed into 34 frames. Reconstruction used the HD-PET algorithm with 4 iterations, 21 subsets, and a 2 mm Full Width at Half Maximum (FWHM) Gaussian filter. MR-based attenuation correction employed the vendor’s dual-echo Ultrashort Echo Time (UTE) sequence, segmenting air, soft tissue, and bone to generate a four-class μ-map for attenuation and scatter correction. If UTE-based MR-based Attenuation Correction (MRAC) failed, the Brain-HiRes protocol was used. All μ-maps were visually inspected prior to use.

All dynamic PET images were motion-corrected and coregistered to each subject’s T1- weighted 3T MRI using SPM12. Parametric images of distribution volume ratio (DVR) were generated using the Multilinear Reference Tissue Model in QModeling (version 3, Matlab R2024b). The inferior cerebellum served as the low-binding reference region, and the globus pallidus as a high-binding region. Mean DVR values were extracted from an a priori meta-ROI of AD-signature region, including entorhinal cortex, amygdala, parahippocampal cortex, fusiform gyrus, and inferior and middle temporal gyri and an RSC ROI using the Desikan-Killiany atlas.

### 3.10. Statistical analysis

Statistical analysis was conducted using the CONN toolbox and R^163^ version 4.4.1 using RStudio, version 2024.04.2^164^. Figures were created with the packages ggplot^165^ and ggseg^166^. The R code used for analyses is openly available (https://github.com/fislarissa/7T_MTL_rsFC). For linear models, homoscedasticity was assessed using the Breusch–Pagan test, and multicollinearity was not evident, with all variance inflation factors (VIFs) below 5. Further, a normal distribution of residuals was assessed using the Shapiro-Wilk test on the standardized residuals. Age, sex, and education were included as covariates in all models.

In sample A (A^-^T^-^ individuals), we investigated the association of rsFC strength and graph measures with age, the interaction of age by *APOE4* group, and the interaction of age by GFAP level. In sample B (AT-Term at baseline and change) and sample C (temporal-lobe and RSC tau PET burden at baseline), we investigated the association of rsFC strength and graph measures with the respective pathology measure, the interaction of pathology by *APOE4* group, and the interaction of pathology by GFAP level. For connections that were part of a significant cluster, standardized β for regressions with the corresponding 95% confidence intervals (95% CI) were calculated. For significant clusters, 95% confidence intervals for the (FDR-corrected) cluster *p*- values were obtained via permutation testing (95% CI*_p_*) to assess the uncertainty due to the finite number of permutations^167^.

Regarding episodic memory performance, we defined in our preregistration^59^ to only assess the connection showing the strongest effect in the respective analysis to avoid multiple testing and to prioritise regional specificity. Specifically, in sample A (A^-^T^-^ individuals), we investigated the association of rsFC strength and the interactions of rsFC by age, rsFC by *APOE4*, and rsFC by GFAP for left BA36 - left dentate gyrus body (connection with the strongest effect for age) and left deep CA3 body - left superficial CA3 body (connection with the strongest effect for age by *APOE4*) with performance. In sample B, we first investigated the relationship of the AT-

Term and episodic memory (at baseline and change). If this relationship was present, we investigated rsFC strength for right BA 36 - left CA1 head (connection with the strongest effect in the AT-Term model) as a moderator, consistent with the resilience and reserve framework^40^. If this relationship was not present, we investigated rsFC as additional predictor in the model, instead of as moderator, as well as the interaction of rsFC by *APOE4* group, and rsFC by GFAP on episodic memory. In sample C, we followed the same procedure regarding MTL and RSC tau and the connections with the strongest effect in the tau models, left CA1 head - left subiculum head and right tail (CA1) - left superficial RSC, respectively.

### 3.11. Control analysis

As planned control analyses^59^, we first included white-matter binding of the tau-PET tracer (global white-matter DVR) and the influence of hippocampal volume (bilateral volume via ASHS segmentation) in in the models to assess off-target tracer binding and early atrophy. We expect no change in our observed associations.

Second, we assessed the specificity of associations of episodic memory, pathology, and rsFC. We therefore investigated the relationship of attention and processing speed via the SDMT and global cognition via CAG instead of the episodic memory composite with pathology and rsFC. We expect no associations of SDMT and CAG with pathology and rsFC.

Third, we assessed whether we could replicate the findings from the resting-state data within our sample, using the breathing-task data (with task effects regressed out). We investigated the connections where we observed effects in the resting-state data and additionally conducted the same analyses as described above including cluster correction. Even though test– retest reliability of rsFC, particularly involving medial temporal lobe regions, is only moderate^119^, motion may be increased during the breathing task due to the nature of the paradigm^136^, and although network-based statistics^167^ introduces an additional source of variability through the cluster-forming threshold, we nevertheless expected to observe broadly similar connectivity patterns.

## List of abbreviations

AD: Alzheimer’s disease
Aβ: Amyloid-beta
*APOE4*: Apolipoprotein E4
ASHS: Automated segmentation of hippocampal subfields
AT-Term: Amyloid-tau composite score (1/(Aβ1-42/Aβ1-40) × p-tau217)
BOLD: Blood-oxygen-level-dependent
CAG: Cognitive age gap
CERAD: Consortium to Establish a Registry for Alzheimer’s Disease
DVR: Distribution volume ratio
DZNE: German Center for Neurodegenerative Diseases
EPI: Echo-planar imaging
FDR: False discovery rate
fMRI: Functional magnetic resonance imaging
GFAP: Glial fibrillary acidic protein
MCI: Mild cognitive impairment
MTL: Medial temporal lobe
NBS: Network-based statistics
PET: Positron emission tomography
p-tau217: Phosphorylated tau 217
RBANS: Repeatable Battery for Assessment of Neuropsychological Status
RCFT: Rey Complex Figure Test and Recognition Trial
ROI: Region of interest
rsFC: Resting-state functional connectivity
SDMT: Symbol Digit Modalities Test
VIF: Variance inflation factor
VLMT: Verbal Learning and Memory Test
WMS: Wechsler Memory Scale

## Declarations

### Ethics approval and consent to participate

All study procedures and experimental protocols were approved by the local Ethics committee of the Medical Faculty, Otto-von-Guericke University Magdeburg (200/19) in accordance with the Declaration of Helsinki. All participants provided written informed consent.

### Consent for publication

Not applicable.

### Availability of data and materials

The datasets generated and/or analyzed during the current study are not publicly available due to the inclusion of sensitive participant information and privacy concerns but are available from the corresponding author on reasonable request. Code for statistical analysis is available at https://github.com/fislarissa/7T_MTL_rsFC.

### Authors’ contributions

Conceptualisation: L.F., S.R.-C., A.M. Methodology: L.F., H.G., A.M. Formal analysis: L.F. Data Acquisition: N.V., B.G.-G., B.S.-W., N.B., L.F. Image processing: L.F., J.H.-F. Visualisation: L.F. Image analysis and modelling: L.F. Investigation: L.F. Supervision: A.M. Funding acquisition: A.M., E.K., S.S., M.K., E.D. Resources: A.M. Writing original draft preparation: LF. Writing – review and editing: All authors. All authors read and approved the final manuscript.

## Acknowledgements

We are grateful to all participants for their time and dedication to this study. We further thank Kathrin Baldauf, Ines Bodewald, Stefanie Hildebrandt, Niklas Kelling, Cindy Lübeck, Hendrik Mattern, and Peter Schulze for their assistance and support regarding the MRI and PET scanning. We also would like to thank the medical-technical assistants (MTAs), in particular Iris Mann, Ulrike Pankratz, and Vivica Sommerfeld as well as our student assistants for their valuable contributions to this study.

## References

1. Ranganath, C. & Ritchey, M. Two cortical systems for memory-guided behaviour. Nat. Rev. Neurosci. 13, 713–726 (2012).

2. Moscovitch, M., Cabeza, R., Winocur, G. & Nadel, L. Episodic Memory and Beyond: The Hippocampus and Neocortex in Transformation. Annu. Rev. Psychol. 67, 105–134 (2016).

3. Braak, H. & Braak, E. Neuropathological stageing of Alzheimer-related changes. Acta Neuropathol. 82, 239–259 (1991).

4. Thal, D. R., Rüb, U., Orantes, M. & Braak, H. Phases of A beta-deposition in the human brain and its relevance for the development of AD. Neurology 58, 1791–1800 (2002).

5. Moscoso, A. et al. Frequency and clinical outcomes associated with tau positron emission tomography positivity. JAMA 334, 229–242 (2025).

6. Jansen, W. J. et al. Prevalence of cerebral amyloid pathology in persons without dementia: A meta-analysis. JAMA 313, 1924 (2015).

7. Ossenkoppele, R. et al. Amyloid and tau PET-positive cognitively unimpaired individuals are at high risk for future cognitive decline. Nat Med 28, 2381–2387 (2022).

8. Spires-Jones, T. L., Attems, J. & Thal, D. R. Interactions of pathological proteins in neurodegenerative diseases. Acta Neuropathol. 134, 187–205 (2017).

9. López-Otín, C., Blasco, M. A., Partridge, L., Serrano, M. & Kroemer, G. Hallmarks of aging: An expanding universe. Cell 186, 243–278 (2023).

10. Suzuki, W. A. & Amaral, D. G. Perirhinal and parahippocampal cortices of the macaque monkey: cortical afferents. J. Comp. Neurol. 350, 497–533 (1994).

11. Amaral, D. & Lavenex, P. Hippocampal Neuroanatomy. in OUP Academic (Oxford University Press, 2006).

12. Valero, M. & de la Prida, L. M. The hippocampus in depth: a sublayer-specific perspective of entorhinal-hippocampal function. Curr Opin Neurobiol 52, 107–114 (2018).

13. Kobayashi, Y. & Amaral, D. G. Macaque monkey retrosplenial cortex: II. Cortical afferents. J Comp Neurol 466, 48–79 (2003).

14. Yamawaki, N. et al. Long-range inhibitory intersection of a retrosplenial thalamocortical circuit by apical tuft-targeting CA1 neurons. Nat Neurosci 22, 618–626 (2019).

15. Braak, H., Alafuzoff, I., Arzberger, T., Kretzschmar, H. & Del Tredici, K. Staging of Alzheimer disease-associated neurofibrillary pathology using paraffin sections and immunocytochemistry. Acta Neuropathol. 112, 389–404 (2006).

16. Ben-Nejma, I. R. H. et al. Increased soluble amyloid-beta causes early aberrant brain network hypersynchronisation in a mature-onset mouse model of amyloidosis. Acta Neuropathol. Commun. 7, 180 (2019).

17. Roemer-Cassiano, S. N. et al. Amyloid-associated hyperconnectivity drives tau spread across connected brain regions in Alzheimer’s disease. Science Translational Medicine (2025) doi:10.1126/scitranslmed.adp2564.

18. Palop, J. J. et al. Aberrant excitatory neuronal activity and compensatory remodeling of inhibitory hippocampal circuits in mouse models of Alzheimer’s disease. Neuron 55, 697–711 (2007).

19. Ziontz, J., Adams, J. N., Harrison, T. M., Baker, S. L. & Jagust, W. J. Hippocampal Connectivity with Retrosplenial Cortex is Linked to Neocortical Tau Accumulation and Memory Function. J. Neurosci. 41, 8839–8847 (2021).

20. Corriveau-Lecavalier, N., Adams, J. N., Fischer, L., Molloy, E. N. & Maass, A. Cerebral hyperactivation across the Alzheimer’s disease pathological cascade. Brain Commun. 6, fcae376 (2024).

21. Mayeux, R. Epidemiology of neurodegeneration. Annu. Rev. Neurosci. 26, 81–104 (2003).

22. Liu, C.-C., Liu, C.-C., Kanekiyo, T., Xu, H. & Bu, G. Apolipoprotein E and Alzheimer disease: risk, mechanisms and therapy. Nat. Rev. Neurol. 9, 106–118 (2013).

23. Selkoe, D. J. & Hardy, J. The amyloid hypothesis of Alzheimer’s disease at 25 years. EMBO Mol. Med. 8, 595–608 (2016).

24. Tabuena, D. R. et al. Neuronal APOE4-induced early hippocampal network hyperexcitability in Alzheimer’s disease pathogenesis. Nat Aging (2026) doi:10.1038/s43587-026-01096-0.

25. Escartin, C. et al. Reactive astrocyte nomenclature, definitions, and future directions. Nat. Neurosci. 24, 312–325 (2021).

26. Kim, H. Y. et al. The rise of astrocytes: are they guardians or troublemakers of the brain disorder? Exp Mol Med 58, 301–318 (2026).

27. Zott, B. & Konnerth, A. Impairments of glutamatergic synaptic transmission in Alzheimer’s disease. Semin. Cell Dev. Biol. 139, 24–34 (2023).

28. Fischer, L. et al. Differential effects of aging, Alzheimer’s pathology, and APOE4 on longitudinal functional connectivity and episodic memory in older adults. Alzheimers. Res. Ther. **17**, 91 (2025).

29. Hrybouski, S. et al. Aging and Alzheimer’s disease have dissociable effects on local and regional medial temporal lobe connectivity. Brain Commun 5, fcad245 (2023).

30. Cassady, K. E. et al. Alzheimer’s Pathology Is Associated with Dedifferentiation of Intrinsic Functional Memory Networks in Aging. Cereb Cortex 31, 4781–4793 (2021).

31. Koen, J. D., Hauck, N. & Rugg, M. D. The Relationship between Age, Neural Differentiation, and Memory Performance. J. Neurosci. 39, 149–162 (2019).

32. Reagh, Z. M. et al. Functional imbalance of anterolateral entorhinal cortex and hippocampal dentate/CA3 underlies age-related object pattern separation deficits. Neuron 97, 1187–1198.e4 (2018).

33. Wilson, I. A., Ikonen, S., Gallagher, M., Eichenbaum, H. & Tanila, H. Age-associated alterations of hippocampal place cells are subregion specific. J Neurosci 25, 6877–6886 (2005).

34. Haberman, R. P., Colantuoni, C., Koh, M. T. & Gallagher, M. Behaviorally activated mRNA expression profiles produce signatures of learning and enhanced inhibition in aged rats with preserved memory. PLoS One 8, e83674 (2013).

35. Koen, J. D. & Rugg, M. D. Neural dedifferentiation in the aging brain. Trends Cogn. Sci. 23, 547– 559 (2019).

36. Adams, J. N. et al. Entorhinal–Hippocampal Circuit Integrity Is Related to Mnemonic Discrimination and Amyloid-β Pathology in Older Adults. J. Neurosci. 42, 8742–8753 (2022).

37. Adams, J. N. et al. Functional network structure supports resilience to memory deficits in cognitively normal older adults with amyloid-β pathology. Sci. Rep. 13, 13953 (2023).

38. Elman, J. A. et al. Neural compensation in older people with brain amyloid-β deposition. Nat. Neurosci. 17, 1316–1318 (2014).

39. Knights, E., Henson, R. N., Morcom, A., Mitchell, D. J. & Tsvetanov, K. A. Neural evidence of functional compensation for fluid intelligence in healthy ageing. Elife 13, RP93327 (2025).

40. Stern, Y. et al. Whitepaper: Defining and investigating cognitive reserve, brain reserve, and brain maintenance. Alzheimers Dement 16, 1305–1311 (2020).

41. Adams, J. N., Maass, A., Harrison, T. M., Baker, S. L. & Jagust, W. J. Cortical tau deposition follows patterns of entorhinal functional connectivity in aging. Elife 8, (2019).

42. Franzmeier, N. et al. Functional brain architecture is associated with the rate of tau accumulation in Alzheimer’s disease. Nat. Commun. 11, 347 (2020).

43. Pooler, A. M., Phillips, E. C., Lau, D. H. W., Noble, W. & Hanger, D. P. Physiological release of endogenous tau is stimulated by neuronal activity. EMBO Rep 14, 389–394 (2013).

44. Wu, J. W. et al. Neuronal activity enhances tau propagation and tau pathology in vivo. Nat. Neurosci. 19, 1085–1092 (2016).

45. Igarashi, K. M. Entorhinal cortex dysfunction in Alzheimer’s disease. Trends in Neurosciences 46, 124–136 (2023).

46. Scharfman, H. E. & Chao, M. V. The entorhinal cortex and neurotrophin signaling in Alzheimer’s disease and other disorders. Cogn Neurosci 4, 123–135 (2013).

47. Braak, H. & Del Tredici, K. Spreading of Tau Pathology in Sporadic Alzheimer’s Disease Along Cortico-cortical Top-Down Connections. Cereb Cortex 28, 3372–3384 (2018).

48. Fischer, L. et al. Longitudinal functional connectivity during rest and task is differentially related to Alzheimer’s pathology and episodic memory in older adults. Sci Rep 15, 38499 (2025).

49. Göschel, L. et al. Plasma p-tau181 and GFAP reflect 7T MR-derived changes in Alzheimer’s disease: A longitudinal study of structural and functional MRI and MRS. Alzheimers Dement 20, 8684–8699 (2024).

50. Maass, A., Berron, D., Libby, L. A., Ranganath, C. & Düzel, E. Functional subregions of the human entorhinal cortex. Elife 4, e06426 (2015).

51. Grande, X., Sauvage, M. M., Becke, A., Düzel, E. & Berron, D. Transversal functional connectivity and scene-specific processing in the human entorhinal-hippocampal circuitry. Elife 11, e76479 (2022).

52. Reznik, D., Margulies, D. S., Witter, M. P. & Doeller, C. F. Evidence for convergence of distributed cortical processing in band-like functional zones in human entorhinal cortex. Curr. Biol. 34, 5457– 5469.e2 (2024).

53. Zhang, K. et al. Differential Laminar Activation Dissociates Encoding and Retrieval in the Human Medial and Lateral Entorhinal Cortex. J. Neurosci. 43, 2874–2884 (2023).

54. Pfaffenrot, V. et al. Characterizing BOLD activation patterns in the human hippocampus with laminar fMRI. *Imaging neuroscience (Cambridge*, Mass.) 3, (2025).

55. Ahmadi, K. et al. Blood volume-sensitive laminar fMRI with VASO in human hippocampus: Capabilities and biophysical challenges at clinical 7T scanners. Imaging Neurosci. (Camb.) 4, IMAG.a.1197 (2026).

56. Maass, A. et al. Laminar activity in the hippocampus and entorhinal cortex related to novelty and episodic encoding. Nat. Commun. 5, 5547 (2014).

57. Liu, P. et al. Layer-specific changes in sensory cortex across the lifespan in mice and humans. Nature Neuroscience 1–12 (2025).

58. Doehler, J. et al. The 3D structural architecture of the human hand area is nontopographic. J. Neurosci. 43, 3456–3476 (2023).

59. Fischer, L. & Maass, A. Preregistration: Medial temporal lobe functional connectivity at 7 Tesla with aging and Alzheimer’s disease pathology in cognitively unimpaired older adults. https://osf.io/ysr2h/overview (2025) doi:10.17605/OSF.IO/YSR2H.

60. Behrenbruch, N. et al. A physically and mentally active lifestyle relates to younger brain and cognitive age. GeroScience 1–21 (2025).

61. Morgado, B. et al. Assessment of immunoprecipitation with subsequent immunoassays for the blood-based diagnosis of Alzheimer’s disease. European Archives of Psychiatry and Clinical Neuroscience 275, 2215–2227 (2024).

62. Maass, A. et al. Associations of [F]PI-2620 Binding with Memory and Phosphorylated Tau 217 in Cognitively Unimpaired Older Adults. J Nucl Med (2026) doi:10.2967/jnumed.125.271927.

63. Gatzen, J. et al. Episodic memory network connectivity with aging and Alzheimer’s disease pathology in cognitively unimpaired older adults. Alzheimers. Dement. 21 **Suppl 7**, e108678 (2025).

64. Feczko, E., Augustinack, J. C., Fischl, B. & Dickerson, B. C. An MRI-based method for measuring volume, thickness and surface area of entorhinal, perirhinal, and posterior parahippocampal cortex. Neurobiol Aging 30, 420–431 (2009).

65. Dickerson, B. C. et al. Differential effects of aging and Alzheimer’s disease on medial temporal lobe cortical thickness and surface area. Neurobiol Aging 30, 432–440 (2009).

66. Palomero-Gallagher, N., Kedo, O., Mohlberg, H., Zilles, K. & Amunts, K. Multimodal mapping and analysis of the cyto- and receptorarchitecture of the human hippocampus. Brain Struct Funct 225, 881–907 (2020).

67. Cabeza, R. et al. Maintenance, reserve and compensation: the cognitive neuroscience of healthy ageing. Nat. Rev. Neurosci. 19, 701–710 (2018).

68. Honey, C. J. et al. Predicting human resting-state functional connectivity from structural connectivity. Proceedings of the National Academy of Sciences 106, 2035–2040 (2009).

69. Honey, C. J., Thivierge, J.-P. & Sporns, O. Can structure predict function in the human brain? Neuroimage 52, 766–776 (2010).

70. Liu, Z.-Q., Betzel, R. F. & Misic, B. Benchmarking functional connectivity by the structure and geometry of the human brain. Netw. Neurosci. 6, 937–949 (2022).

71. Insausti, R. & Amaral, D. G. Hippocampal Formation. in The Human Nervous System 871–914 (Elsevier, 2004).

72. Andersen, P., Morris, R., Amaral, D., Bliss, T. & O’Keefe, J. The Hippocampus Book. (Oxford University Press, 2006).

73. Cholvin, T. & Bartos, M. The dentate gyrus efficiently converges LEC and MEC inputs into multimodal, highly specific and reliable environmental representations. Nat Neurosci (2026) doi:10.1038/s41593-026-02240-0.

74. Chauveau, L. et al. Anterior-temporal network hyperconnectivity is key to Alzheimer’s disease: from ageing to dementia. Brain (2025) doi:10.1093/brain/awaf008.

75. Damoiseaux, J. S., Viviano, R. P., Yuan, P. & Raz, N. Differential effect of age on posterior and anterior hippocampal functional connectivity. Neuroimage 133, 468–476 (2016).

76. Dalton, M. A., McCormick, C., De Luca, F., Clark, I. A. & Maguire, E. A. Functional connectivity along the anterior-posterior axis of hippocampal subfields in the ageing human brain. Hippocampus 29, 1049–1062 (2019).

77. Adler, D. H. et al. Characterizing the human hippocampus in aging and Alzheimer’s disease using a computational atlas derived from ex vivo MRI and histology. Proc. Natl. Acad. Sci. U. S. A. 115, 4252–4257 (2018).

78. Bobinski, M. et al. Neuronal and volume loss in CA1 of the hippocampal formation uniquely predicts duration and severity of Alzheimer disease. Brain Res. 805, 267–269 (1998).

79. Small, S. A., Chawla, M. K., Buonocore, M., Rapp, P. R. & Barnes, C. A. Imaging correlates of brain function in monkeys and rats isolates a hippocampal subregion differentially vulnerable to aging. Proc Natl Acad Sci U S A 101, 7181–7186 (2004).

80. Achard, S. & Bullmore, E. Efficiency and cost of economical brain functional networks. PLoS Comput Biol 3, e17 (2007).

81. Yassa, M. A. et al. Pattern separation deficits associated with increased hippocampal CA3 and dentate gyrus activity in nondemented older adults. Hippocampus 21, 968–979 (2011).

82. Sinha, N. et al. APOE ε4 status in healthy older African Americans is associated with deficits in pattern separation and hippocampal hyperactivation. Neurobiol. Aging 69, 221–229 (2018).

83. Patrylo, P. R. & Williamson, A. The effects of aging on dentate circuitry and function. Prog Brain Res 163, 679–696 (2007).

84. Wilson, I. A., Gallagher, M., Eichenbaum, H. & Tanila, H. Neurocognitive aging: prior memories hinder new hippocampal encoding. Trends Neurosci 29, 662–670 (2006).

85. Leal, S. L. & Yassa, M. A. Perturbations of neural circuitry in aging, mild cognitive impairment, and Alzheimer’s disease. Ageing Res. Rev. 12, 823–831 (2013).

86. Ashton, N. J. et al. Diagnostic Accuracy of a Plasma Phosphorylated Tau 217 Immunoassay for Alzheimer Disease Pathology. JAMA Neurol 81, 255–263 (2024).

87. Morgado, B. et al. Evaluation of a fully automated assay for the measurement of plasma pTau217 and a composite score integrating the ratio Aβ1-42/1-40 as biomarkers of Alzheimer’s disease. Eur. Arch. Psychiatry Clin. Neurosci. (2025) doi:10.1007/s00406-025-02123-8.

88. Verret, L. et al. Inhibitory interneuron deficit links altered network activity and cognitive dysfunction in Alzheimer model. Cell 149, 708–721 (2012).

89. Zott, B. et al. A vicious cycle of β amyloid–dependent neuronal hyperactivation. Science (2019) doi:10.1126/science.aay0198.

90. Busche, M. A. et al. Clusters of hyperactive neurons near amyloid plaques in a mouse model of Alzheimer’s disease. Science 321, 1686–1689 (2008).

91. Ellingford, R. et al. Alzheimer’s disease pathology degrades an NMDA receptor-dependent spontaneous activity pattern in cortico-hippocampal circuits. Neuron (2026) doi:10.1016/j.neuron.2026.02.027.

92. Šišková, Z. et al. Dendritic structural degeneration is functionally linked to cellular hyperexcitability in a mouse model of Alzheimer’s disease. Neuron 84, 1023–1033 (2014).

93. Thangavel, R., Van Hoesen, G. W. & Zaheer, A. The abnormally phosphorylated tau lesion of early Alzheimer’s disease. Neurochem Res 34, 118–123 (2009).

94. Braak, E. & Braak, H. Alzheimer’s disease: transiently developing dendritic changes in pyramidal cells of sector CA1 of the Ammon’s horn. Acta Neuropathol. 93, 323–325 (1997).

95. Kaufman, S. K., Del Tredici, K., Thomas, T. L., Braak, H. & Diamond, M. I. Tau seeding activity begins in the transentorhinal/entorhinal regions and anticipates phospho-tau pathology in Alzheimer’s disease and PART. Acta Neuropathol 136, 57–67 (2018).

96. Ahnaou, A. et al. Emergence of early alterations in network oscillations and functional connectivity in a tau seeding mouse model of Alzheimer’s disease pathology. Sci. Rep. 7, 14189 (2017).

97. Ji, C. et al. Neuronal hypofunction and network dysfunction in a mouse model at an early stage of tauopathy. Alzheimers. Dement. 20, 7954–7970 (2024).

98. Harris, S. S. et al. Alzheimer’s disease patient-derived high-molecular-weight tau impairs bursting in hippocampal neurons. Cell 188, 3775–3788.e21 (2025).

99. Insausti, R., Amaral, D. G. & Cowan, W. M. The entorhinal cortex of the monkey: II. Cortical afferents. J. Comp. Neurol. 264, 356–395 (1987).

100. Delpech, J.-C. et al. Wolframin-1-expressing neurons in the entorhinal cortex propagate tau to CA1 neurons and impair hippocampal memory in mice. Sci Transl Med 13, eabe8455 (2021).

101. Ferrari-Souza, J. P. et al. Microglia modulate Aβ-dependent astrocyte reactivity in Alzheimer’s disease. Nat Neurosci 29, 81–87 (2026).

102. Varma, V. R. et al. Longitudinal progression of blood biomarkers reveals a key role of reactive astrocytosis in preclinical Alzheimer’s disease. Med 6, 100724 (2025).

103. Pereira, J. B. et al. Plasma GFAP is an early marker of amyloid-β but not tau pathology in Alzheimer’s disease. Brain 144, 3505–3516 (2021).

104. Gobbo, F. et al. Astrocyte Proximity Protects Synapses From Human Amyloid-Beta Induced Degeneration in a Mouse Ex Vivo Model of Early Alzheimer’s Disease. European Journal of Neuroscience 63, e70480 (2026).

105. Chiotis, K. et al. Tracking reactive astrogliosis in autosomal dominant and sporadic Alzheimer’s disease with multi-modal PET and plasma GFAP. Mol Neurodegener 18, 60 (2023).

106. Youn, W., Yun, M., Lee, C. J. & Schöll, M. Cautions on utilizing plasma GFAP level as a biomarker for reactive astrocytes in neurodegenerative diseases. Molecular Neurodegeneration 20, 54 (2025).

107. Sánchez-Juan, P. et al. Serum GFAP levels correlate with astrocyte reactivity, post-mortem brain atrophy and neurofibrillary tangles. Brain 147, 1667–1679 (2024).

108. Gogola, A., et al. Complex interplay of astrogliosis and pathology in preclinical Alzheimer’s disease. Alzheimer’s & Dementia 22, e71520 (2026).

109. Gonzalez, J. et al. Subspace communication in the hippocampal-retrosplenial axis. Nature (2026) doi:10.1038/s41586-026-10481-z.

110. Pearson, R. C., Esiri, M. M., Hiorns, R. W., Wilcock, G. K. & Powell, T. P. Anatomical correlates of the distribution of the pathological changes in the neocortex in Alzheimer disease. Proc Natl Acad Sci U S A 82, 4531–4534 (1985).

111. Huber, L. et al. High-resolution CBV-fMRI allows mapping of laminar activity and connectivity of cortical input and output in human M1. Neuron 96, 1253–1263.e7 (2017).

112. Harrison, T. M. et al. Tau deposition is associated with functional isolation of the hippocampus in aging. Nat. Commun. 10, 4900 (2019).

113. Vogel, J. W. et al. Connectome-based modelling of neurodegenerative diseases: towards precision medicine and mechanistic insight. Nat. Rev. Neurosci. 24, 620–639 (2023).

114. Fonseca, C. S. et al. Tau accumulation and atrophy predict amyloid independent cognitive decline in aging. Alzheimers Dement 20, 2526–2537 (2024).

115. Stroh, A., Schweiger, S., Ramirez, J.-M. & Tüscher, O. The selfish network: how the brain preserves behavioral function through shifts in neuronal network state. Trends Neurosci 47, 246–258 (2024).

116. Giorgio, J., Adams, J. N., Maass, A., Jagust, W. J. & Breakspear, M. Amyloid induced hyperexcitability in default mode network drives medial temporal hyperactivity and early tau accumulation. Neuron 112, 676–686.e4 (2024).

117. Diersch, N., Valdes-Herrera, J. P., Tempelmann, C. & Wolbers, T. Increased Hippocampal Excitability and Altered Learning Dynamics Mediate Cognitive Mapping Deficits in Human Aging. J Neurosci 41, 3204–3221 (2021).

118. Schott, B. H. et al. Inhibitory temporo-parietal effective connectivity is associated with explicit memory performance in older adults. iScience 26, 107765 (2023).

119. Noble, S., Scheinost, D. & Constable, R. T. A decade of test-retest reliability of functional connectivity: A systematic review and meta-analysis. Neuroimage 203, 116157 (2019).

120. Birn, R. M. et al. The effect of scan length on the reliability of resting-state fMRI connectivity estimates. Neuroimage 83, 550–558 (2013).

121. De Martino, F. et al. Whole brain high-resolution functional imaging at ultra high magnetic fields: an application to the analysis of resting state networks. Neuroimage 57, 1031–1044 (2011).

122. Huber, L. R. et al. LayNii: A software suite for layer-fMRI. Neuroimage 237, 118091 (2021).

123. Tsvetanov, K. A., Henson, R. N. A. & Rowe, J. B. Separating vascular and neuronal effects of age on fMRI BOLD signals. Philos Trans R Soc Lond B Biol Sci 376, 20190631 (2020).

124. Kucyi, A. et al. Intracranial Electrophysiology Reveals Reproducible Intrinsic Functional Connectivity within Human Brain Networks. The Journal of neuroscience : the official journal of the Society for Neuroscience 38, (2018).

125. Ma, Z., Zhang, Q., Tu, W. & Zhang, N. Gaining insight into the neural basis of resting-state fMRI signal. Neuroimage 250, 118960 (2022).

126. Vafaii, H. et al. Multimodal measures of spontaneous brain activity reveal both common and divergent patterns of cortical functional organization. Nature Communications 15, 229 (2024).

127. Vizioli, L. et al. Lowering the thermal noise barrier in functional brain mapping with magnetic resonance imaging. Nat Commun 12, 5181 (2021).

128. Ehrensperger, M. M., Berres, M., Taylor, K. I. & Monsch, A. U. Early detection of Alzheimer’s disease with a total score of the German CERAD. J Int Neuropsychol Soc 16, 910–920 (2010).

129. Randolph, C., Tierney, M. C., Mohr, E. & Chase, T. N. The Repeatable Battery for the Assessment of Neuropsychological Status (RBANS): preliminary clinical validity. J. Clin. Exp. Neuropsychol. 20, 310–319 (1998).

130. Helmstaedter, C. & Durwen, H. F. The Verbal Learning and Retention Test. A useful and differentiated tool in evaluating verbal memory performance. Schweizer Archiv fur Neurologie und Psychiatrie 141, (1990).

131. Meyers, J. E. & Meyers, K. R. Rey complex figure test under four different administration procedures. The Clinical Neuropsychologist 63–67 (2007).

132. Wechsler, D. WMS-R: Wechsler Memory Scale--Revised : Manual. (1987).

133. Smith, A. Symbol Digit Modalities Test: Manual. (2002).

134. Pezzoli, S. et al. Successful cognitive aging is associated with thicker anterior cingulate cortex and lower tau deposition compared to typical aging. Alzheimers. Dement. 20, 341–355 (2024).

135. Beheshti, I., Nugent, S., Potvin, O. & Duchesne, S. Bias-adjustment in neuroimaging-based brain age frameworks: A robust scheme. NeuroImage Clin. 24, 102063 (2019).

136. Liu, P. et al. Cerebrovascular reactivity mapping using intermittent breath modulation. Neuroimage 215, 116787 (2020).

137. Functional Imaging Laboratory UCL. Statistical Parametric Mapping. Statistical Parametric Mapping https://www.fil.ion.ucl.ac.uk/spm/ (2021).

138. Henschel, L., Kügler, D. & Reuter, M. FastSurferVINN: Building resolution-independence into deep learning segmentation methods-A solution for HighRes brain MRI. Neuroimage 251, 118933 (2022).

139. Henschel, L. et al. FastSurfer - A fast and accurate deep learning based neuroimaging pipeline. Neuroimage 219, 117012 (2020).

140. Fischl, B. FreeSurfer. Neuroimage 62, 774–781 (2012).

141. Berron, D. et al. A protocol for manual segmentation of medial temporal lobe subregions in 7 Tesla MRI. NeuroImage Clin. 15, 466–482 (2017).

142. Yushkevich, P. A. et al. Quantitative comparison of 21 protocols for labeling hippocampal subfields and parahippocampal subregions in in vivo MRI: towards a harmonized segmentation protocol. Neuroimage 111, 526–541 (2015).

143. Xie, L. et al. Baseline structural MRI and plasma biomarkers predict longitudinal structural atrophy and cognitive decline in early Alzheimer’s disease. Alzheimers. Res. Ther. 15, 79 (2023).

144. Xie, L. et al. Automated segmentation of medial temporal lobe subregions on in vivo T1- weighted MRI in early stages of Alzheimer’s disease. Hum. Brain Mapp. 40, 3431–3451 (2019).

145. Canada, K. L. et al. Development and validation of a quality control procedure for automatic segmentation of hippocampal subfields. Hippocampus 33, 1048–1057 (2023).

146. Raz, N., Daugherty, A. M., Bender, A. R., Dahle, C. L. & Land, S. Volume of the hippocampal subfields in healthy adults: differential associations with age and a pro-inflammatory genetic variant. Brain Structure and Function 220, 2663–2674 (2014).

147. Gellersen, H. M. et al. Medial temporal lobe structure, mnemonic and perceptual discrimination in healthy older adults and those at risk for mild cognitive impairment. Neurobiol Aging 122, 88–106 (2023).

148. Gellersen, H., Naspi, L., Przygodda, X., Düzel, E. & Berron, D. Dynamic shifts between pattern separation and reinstatement in human CA3 and dentate gyrus. bioRxiv (2026) doi:10.64898/2026.04.24.720552.

149. Masharipov, R., Knyazeva, I., Korotkov, A., Cherednichenko, D. & Kireev, M. Comparison of whole-brain task-modulated functional connectivity methods for fMRI task connectomics. Commun. Biol. 7, 1402 (2024).

150. Whitfield-Gabrieli, S. NITRC: Artifact detection tools (ART): Tool/resource info. NITRC https://www.nitrc.org/projects/artifact_detect/ (2015).

151. Nieto-Castanon, A. Handbook of Functional Connectivity Magnetic Resonance Imaging Methods in CONN. (Hilbert Press, 2020).

152. Behzadi, Y., Restom, K., Liau, J. & Liu, T. T. A component based noise correction method (CompCor) for BOLD and perfusion based fMRI. Neuroimage 37, 90–101 (2007).

153. Muschelli, J. et al. Reduction of motion-related artifacts in resting state fMRI using aCompCor. Neuroimage 96, 22–35 (2014).

154. Fox, M. D. & Raichle, M. E. Spontaneous fluctuations in brain activity observed with functional magnetic resonance imaging. Nat. Rev. Neurosci. 8, 700–711 (2007).

155. Hayek, D. et al. The In Vivo Microstructural Profile of Human Hippocampal Subfield CA1 and Its Relation to Memory Performance. Hum Brain Mapp 47, e70542 (2026).

156. Huber, L. et al. Layer-dependent functional connectivity methods. Prog. Neurobiol. 207, 101835 (2021).

157. Libby, L. A., Ekstrom, A. D., Ragland, J. D. & Ranganath, C. Differential connectivity of perirhinal and parahippocampal cortices within human hippocampal subregions revealed by high-resolution functional imaging. J. Neurosci. 32, 6550–6560 (2012).

158. Benjamini, Y. & Hochberg, Y. Controlling the false discovery rate: A practical and powerful approach to multiple testing. J. R. Stat. Soc. Series B Stat. Methodol. 57, 289–300 (1995).

159. Bullmore, E. & Sporns, O. Complex brain networks: graph theoretical analysis of structural and functional systems. Nat. Rev. Neurosci. 10, 186–198 (2009).

160. Rubinov, M. & Sporns, O. Complex network measures of brain connectivity: uses and interpretations. Neuroimage 52, 1059–1069 (2010).

161. Latora, V. & Marchiori, M. Efficient behavior of small-world networks. Phys Rev Lett 87, 198701 (2001).

162. Fornito, A., Zalesky, A. & Bullmore, E. Fundamentals of Brain Network Analysis. (Academic Press, San Diego, CA, 2016).

163. R Core Team. R: A language and environment for statistical computing. R Foundation for Statistical Computing. (2024).

164. RStudio Team. RStudio: Integrated Development for R. RStudio. (2024).

165. Wickham, H. *Ggplot2: Elegant Graphics for Data Analysis*. (Springer, New York, NY, 2009).

166. Mowinckel, A. M. & Vidal-Piñeiro, D. Visualization of brain statistics with R packages *ggseg* and *ggseg3d*. Adv. Methods Pract. Psychol. Sci. 3, 466–483 (2020).

167. Zalesky, A., Fornito, A. & Bullmore, E. T. Network-based statistic: identifying differences in brain networks. Neuroimage 53, 1197–1207 (2010).

