## Supplementary Material for "Mesoscale medial temporal lobe connectivity patterns relate to tau pathology and memory in older adults"

### Supplementary Tables

#### Table S1. Sample A: Linear model of effects of age on functional connectivity

| | rsFC between left BA 36 - left dentate gyrus body | | | | | | | |
|---|---|---|---|---|---|---|---|---|
| *Predictors* | *Estimates* | *std. Error* | *std. Beta* | *standardized std. Error* | *CI* | *standardized CI* | *Statistic* | *p* |
| (Intercept) | 0.80 | 0.35 | 0.09 | 0.21 | 0.09 – 1.51 | -0.33 – 0.51 | 2.26 | **0.028** |
| Age in years | -0.02 | 0.00 | -0.46 | 0.13 | -0.03 – -0.01 | -0.71 – -0.20 | -3.63 | **0.001** |
| *APOE4* Group [carrier] | -0.08 | 0.08 | -0.37 | 0.36 | -0.23 – 0.07 | -1.10 – 0.35 | -1.03 | 0.308 |
| Sex [male] | -0.01 | 0.06 | -0.06 | 0.27 | -0.13 – 0.10 | -0.61 – 0.49 | -0.23 | 0.816 |
| Education in years | 0.03 | 0.01 | 0.31 | 0.13 | 0.00 – 0.05 | 0.04 – 0.57 | 2.33 | **0.023** |
| Observations | 57 | | | | | | | |
| $R^2$ / $R^2$ adjusted | 0.248 / 0.190 | | | | | | | |

Functional connectivity in amyloid- and tau-negative older adults was used as dependent variable, age in years at baseline, *APOE4* group, sex, and education in years were used as independent variables. rsFC = resting-state functional connectivity. BA = Brodmann Area. CI = 95% confidence interval.

#### Table S2. Sample A: Linear model of effects of age on functional connectivity

| | rsFC between left BA36 - right deep subiculum body | | | | | | | |
|---|---|---|---|---|---|---|---|---|
| *Predictors* | *Estimates* | *std. Error* | *std. Beta* | *standardized std. Error* | *CI* | *standardized CI* | *Statistic* | *p* |
| (Intercept) | 0.89 | 0.31 | 0.00 | 0.22 | 0.27 – 1.51 | -0.43 – 0.44 | 2.89 | **0.006** |

| | | | | | | | | |
|---|---|---|---|---|---|---|---|---|
| Age in years | -0.01 | 0.00 | -0.46 | 0.13 | -0.02 – -0.01 | -0.72 – -0.21 | -3.61 | **0.001** |
| *APOE4* Group [carrier] | -0.07 | 0.07 | -0.37 | 0.37 | -0.20 – 0.07 | -1.11 – 0.37 | -1.00 | 0.324 |
| Sex [male] | 0.02 | 0.05 | 0.09 | 0.28 | -0.08 – 0.12 | -0.48 – 0.65 | 0.31 | 0.757 |
| Education in years | 0.01 | 0.01 | 0.15 | 0.14 | -0.01 – 0.03 | -0.12 – 0.43 | 1.12 | 0.266 |
| Observations | 57 | | | | | | | |
| $R^2$ / $R^2$ adjusted | 0.210 / 0.149 | | | | | | | |

Functional connectivity in amyloid- and tau-negative older adults was used as dependent variable, age in years at baseline, *APOE4* group, sex, and education in years were used as independent variables. rsFC = resting-state functional connectivity. BA = Brodmann Area. CI = 95% confidence interval.

### Table S3. Sample A: Linear model of effects of age on local efficiency

| | | | | | | | | |
|---|---|---|---|---|---|---|---|---|
| | | | | **Network Local Efficiency** | | | | |
| *Predictors* | *Estimates* | *std. Error* | *std. Beta* | *standardized std. Error* | *CI* | *standardized CI* | *Statistic* | *p* |
| (Intercept) | 0.64 | 0.15 | 0.23 | 0.23 | 0.34 – 0.93 | -0.23 – 0.69 | 4.32 | **<0.001** |
| Age in years | -0.00 | 0.00 | -0.24 | 0.14 | -0.01 – -0.00 | -0.52 – -0.03 | -1.75 | **0.043** |
| *APOE4* Group [carrier] | -0.02 | 0.03 | -0.22 | 0.40 | -0.08 – 0.05 | -1.02 – 0.57 | -0.57 | 0.572 |
| Sex [male] | -0.03 | 0.02 | -0.36 | 0.30 | -0.08 – 0.02 | -0.96 – 0.25 | -1.19 | 0.241 |
| Education in years | 0.01 | 0.01 | 0.23 | 0.14 | -0.00 – 0.02 | -0.06 – 0.52 | 1.60 | 0.115 |
| Observations | 57 | | | | | | | |
| $R^2$ / $R^2$ adjusted | 0.303 / 0.234 | | | | | | | |

The graph measure local efficiency in amyloid- and tau-negative older adults was used as dependent variable, age in years at baseline, *APOE4* group, sex, and education in years were used as independent variables. CI = 95% confidence interval.

### Table S4. Sample A: Linear model of effects of age on local efficiency

| | Left CA3 head local efficiency | | | | | | | |
|---|---|---|---|---|---|---|---|---|
| *Predictors* | *Estimates* | *std. Error* | *std. Beta* | *standardized std. Error* | *CI* | *standardized CI* | *Statistic* | *p* |
| (Intercept) | 2.55 | 0.51 | -0.22 | 0.19 | 1.51 – 3.58 | -0.61 – 0.17 | 4.97 | **<0.001** |
| Age in years | -0.04 | 0.01 | -0.66 | 0.13 | -0.05 – -0.02 | -0.93 – -0.39 | -4.94 | **<0.001** |
| *APOE4* Group [carrier] | 0.13 | 0.12 | 0.42 | 0.37 | -0.11 – 0.38 | -0.34 – 1.18 | 1.12 | 0.268 |
| Sex [male] | 0.10 | 0.08 | 0.32 | 0.26 | -0.07 – 0.27 | -0.21 – 0.85 | 1.21 | 0.234 |
| Education in years | 0.03 | 0.02 | 0.20 | 0.13 | -0.01 – 0.06 | -0.06 – 0.47 | 1.58 | 0.123 |
| Observations | 57 | | | | | | | |
| $R^2$ / $R^2$ adjusted | 0.471 / 0.415 | | | | | | | |

The graph measure local efficiency in amyloid- and tau-negative older adults was used as dependent variable, age in years at baseline, *APOE4* group, sex, and education in years were used as independent variables. CI = 95% confidence interval. *p*-FDR for age was 0.0003.

### Table S5. Sample A: Linear model of interaction effects of age by *APOE4* group on functional connectivity

| | rsFC between left superficial CA3 body - left deep CA3 body | | | | | | | | | |
|---|---|---|---|---|---|---|---|---|---|---|
| *Predictors* | *Estimates* | *std. Error* | *std. Beta* | *standardized std. Error* | *CI* | *standardized CI* | *Statistic* | *std. Statistic* | *p* | *std. p* |
| (Intercept) | 0.80 | 0.36 | -0.12 | 0.22 | 0.09 – 1.52 | -0.56 – 0.31 | 2.25 | -0.57 | **0.028** | 0.568 |
| Age in years | -0.01 | 0.00 | -0.17 | 0.13 | -0.02 – 0.00 | -0.44 – 0.10 | -1.30 | -1.30 | 0.200 | 0.200 |
| *APOE4* Group [carrier] | -3.16 | 0.95 | 0.99 | 0.40 | -5.06 – -1.26 | 0.19 – 1.79 | -3.34 | 2.50 | **0.002** | **0.016** |
| Sex [male] | 0.03 | 0.06 | 0.14 | 0.28 | -0.09 – 0.14 | -0.43 – 0.70 | 0.48 | 0.48 | 0.634 | 0.634 |

| | | | | | | | | | | |
|---|---|---|---|---|---|---|---|---|---|---|
| Education in years | -0.02 | 0.01 | -0.21 | 0.13 | -0.04 – 0.01 | -0.48 – 0.07 | -1.52 | -1.52 | 0.134 | 0.134 |
| Age in years × *APOE4* Group [carrier] | 0.05 | 0.01 | 1.36 | 0.39 | 0.02 – 0.08 | 0.57 – 2.15 | 3.45 | 3.45 | **0.001** | **0.001** |
| Observations | 57 | | | | | | | | | |
| $R^2$ / $R^2$ adjusted | 0.238 / 0.164 | | | | | | | | | |

Functional connectivity in amyloid- and tau-negative older adults was used as dependent variable, age in years at baseline, *APOE4* group, sex, education in years, and the interaction of age by *APOE4* group were used as independent variables. rsFC = resting-state functional connectivity. CI = 95% confidence interval.

### Table S6. Sample A: Linear model of effects of age on functional connectivity in *APOE4* carriers

| | rsFC between left superficial CA3 body - left deep CA3 body | | | | | | | |
|---|---|---|---|---|---|---|---|---|
| *Predictors* | *Estimates* | *std. Error* | *std. Beta* | *standardized std. Error* | *CI* | *standardized CI* | *Statistic* | *p* |
| (Intercept) | -1.00 | 1.15 | -0.36 | 0.28 | -4.20 – 2.20 | -1.15 – 0.43 | -0.87 | 0.433 |
| Age in years | 0.03 | 0.01 | 0.52 | 0.28 | -0.01 – 0.06 | -0.25 – 1.30 | 1.87 | 0.135 |
| Sex [male] | 0.38 | 0.22 | 1.44 | 0.83 | -0.22 – 0.98 | -0.85 – 3.73 | 1.74 | 0.156 |
| Education in years | -0.04 | 0.03 | -0.42 | 0.31 | -0.13 – 0.05 | -1.28 – 0.45 | -1.33 | 0.253 |
| Observations | 8 | | | | | | | |
| $R^2$ / $R^2$ adjusted | 0.654 / 0.462 | | | | | | | |

Functional connectivity in amyloid- and tau-negative older *APOE4* carriers was used as dependent variable, age in years at baseline, sex, and education in years were used as independent variables. rsFC = resting-state functional connectivity. CI = 95% confidence interval.

### Table S7. Sample A: Linear model of effects of age on functional connectivity in *APOE4* non-carriers

| | rsFC between left superficial CA3 body - left deep CA3 body | | | | | | | |
|---|---|---|---|---|---|---|---|---|
| *Predictors* | *Estimates* | *std. Error* | *std. Beta* | *standardized std. Error* | *CI* | *standardized CI* | *Statistic* | *p* |

| | | | | | | | | |
|---|---|---|---|---|---|---|---|---|
| (Intercept) | 0.81 | 0.37 | -0.03 | 0.24 | 0.08 – 1.55 | -0.51 – 0.45 | 2.22 | **0.031** |
| Age in years | -0.01 | 0.00 | -0.18 | 0.14 | -0.02 – 0.00 | -0.47 – 0.11 | -1.22 | 0.229 |
| Sex [male] | 0.01 | 0.06 | 0.05 | 0.31 | -0.11 – 0.13 | -0.58 – 0.68 | 0.16 | 0.873 |
| Education in years | -0.02 | 0.01 | -0.23 | 0.15 | -0.05 – 0.01 | -0.54 – 0.08 | -1.48 | 0.145 |
| Observations | 49 | | | | | | | |
| $R^2$ / $R^2$ adjusted | 0.091 / 0.031 | | | | | | | |

Functional connectivity in amyloid- and tau-negative older *APOE4* non-carriers was used as dependent variable, age in years at baseline, sex, and education in years were used as independent variables. rsFC = resting-state functional connectivity. CI = 95% confidence interval.

### Table S8. Sample A: Linear model of interaction effects of age by *APOE4* group on functional connectivity including all *APOE4* carriers (exploratory analysis)

| | rsFC between left superficial CA3 body - left deep CA3 body | | | | | | | | | |
|---|---|---|---|---|---|---|---|---|---|---|
| *Predictors* | *Estimates* | *std. Error* | *std. Beta* | *standardized std. Error* | *CI* | *standardized CI* | *Statistic* | *std. Statistic* | *p* | *std. p* |
| (Intercept) | 0.81 | 0.36 | -0.23 | 0.22 | 0.09 – 1.53 | -0.66 – 0.20 | 2.26 | -1.07 | **0.028** | 0.289 |
| Age in years | -0.01 | 0.00 | -0.15 | 0.14 | -0.01 – 0.00 | -0.42 – 0.13 | -1.06 | -1.06 | 0.294 | 0.294 |
| *APOE4* Group [carrier] | -2.30 | 0.64 | 0.79 | 0.34 | -3.58 – -1.02 | 0.11 – 1.47 | -3.60 | 2.34 | **0.001** | **0.023** |
| Sex [male] | 0.03 | 0.05 | 0.15 | 0.26 | -0.08 – 0.14 | -0.37 – 0.68 | 0.58 | 0.58 | 0.563 | 0.563 |
| Education in years | -0.02 | 0.01 | -0.23 | 0.13 | -0.04 – 0.00 | -0.48 – 0.03 | -1.75 | -1.75 | 0.086 | 0.086 |
| AT Term | -0.04 | 0.03 | -0.23 | 0.14 | -0.09 – 0.01 | -0.52 – 0.05 | -1.64 | -1.64 | 0.107 | 0.107 |

| | | | | | | | | | | |
|---|---|---|---|---|---|---|---|---|---|---|
| Age in years × *APOE4* Group [carrier] | 0.03 | 0.01 | 1.00 | 0.26 | 0.02 – 0.05 | 0.48 – 1.52 | 3.85 | 3.85 | **<0.001** | **<0.001** |
| Observations | 63 | | | | | | | | | |
| $R^2$ / $R^2$ adjusted | 0.273 / 0.195 | | | | | | | | | |

Functional connectivity in amyloid- and tau-negative older adults and additionally in all 6 APOE4 carriers above the biomarker threshold was used as dependent variable, age in years at baseline, *APOE4* group, sex, education in years, the AT-Term, and the interaction of age by *APOE4* group were used as independent variables. rsFC = resting-state functional connectivity. CI = 95% confidence interval.

### Table S9. Sample A: Linear model of effects of age on functional connectivity in *APOE4* carriers including all *APOE4* carriers (exploratory analysis)

| | **rsFC between left superficial CA3 body - left deep CA3 body** | | | | | | | |
|---|---|---|---|---|---|---|---|---|
| *Predictors* | *Estimates* | *std. Error* | *std. Beta* | *standardized std. Error* | *CI* | *standardized CI* | *Statistic* | *p* |
| (Intercept) | -1.17 | 0.70 | -0.28 | 0.35 | -2.76 – 0.42 | -1.07 – 0.51 | -1.67 | 0.130 |
| Age in years | 0.03 | 0.01 | 0.71 | 0.26 | 0.00 – 0.05 | 0.12 – 1.30 | 2.72 | **0.024** |
| Sex [male] | 0.14 | 0.15 | 0.56 | 0.58 | -0.19 – 0.48 | -0.74 – 1.86 | 0.97 | 0.356 |
| Education in years | -0.03 | 0.03 | -0.30 | 0.28 | -0.10 – 0.04 | -0.95 – 0.34 | -1.07 | 0.314 |
| AT Term | -0.05 | 0.03 | -0.40 | 0.24 | -0.13 – 0.02 | -0.95 – 0.14 | -1.68 | 0.126 |
| Observations | 14 | | | | | | | |
| $R^2$ / $R^2$ adjusted | 0.607 / 0.432 | | | | | | | |

Functional connectivity in amyloid- and tau-negative older *APOE4* carriers and additionally in all 6 APOE4 carriers above the biomarker threshold was used as dependent variable, age in years at baseline, sex, education in years, and the AT-Term were used as independent variables. rsFC = resting-state functional connectivity. CI = 95% confidence interval.

### Table S10. Sample B: Linear model of effects of AT-Term on functional connectivity

| | **rsFC between right BA36 - left CA1 head** | | | | | | | |
|---|---|---|---|---|---|---|---|---|
| *Predictors* | *Estimates* | *std. Error* | *std. Beta* | *standardized std. Error* | *CI* | *standardized CI* | *Statistic* | *p* |

| | | | | | | | | |
|---|---|---|---|---|---|---|---|---|
| (Intercept) | 0.61 | 0.24 | -0.45 | 0.22 | 0.13 – 1.09 | -0.89 – -0.02 | -2.07 | **0.043** |
| AT-Term | 0.13 | 0.03 | 1.13 | 0.29 | 0.06 – 0.20 | 0.55 – 1.71 | 3.91 | **<0.001** |
| Age in years | -0.01 | 0.00 | -0.31 | 0.11 | -0.01 – -0.00 | -0.53 – -0.08 | -2.69 | **0.009** |
| Sex [male] | -0.04 | 0.04 | -0.22 | 0.24 | -0.12 – 0.05 | -0.71 – 0.26 | -0.92 | 0.360 |
| Education in years | 0.00 | 0.01 | 0.03 | 0.12 | -0.02 – 0.02 | -0.21 – 0.27 | 0.25 | 0.802 |
| Observations | 72 | | | | | | | |
| $R^2$ / $R^2$ adjusted | 0.239 / 0.193 | | | | | | | |

Functional connectivity in older adults with plasma biomarkers was used as dependent variable, the AT-Term (1/ ($A\beta_{1-42}$/$A\beta_{1-40}$) * p-$tau_{217}$), age in years at baseline, sex, and education in years were used as independent variables. AT-Term values were log-transformed. rsFC = resting-state functional connectivity. CI = 95% confidence interval.

### Table S11. Sample B: Linear model of effects of change in AT-Term on functional connectivity

| | rsFC between right BA36 - left CA1 head | | | | | | | |
|---|---|---|---|---|---|---|---|---|
| *Predictors* | *Estimates* | *std. Error* | *std. Beta* | *standardized std. Error* | *CI* | *standardized CI* | *Statistic* | *p* |
| (Intercept) | 1.37 | 0.28 | -0.40 | 0.27 | 0.81 – 1.94 | -0.95 – 0.15 | -1.47 | **0.150** |
| Change in AT-Term | 0.09 | 0.02 | 1.18 | 0.34 | 0.05 – 0.14 | 0.49 – 1.86 | 3.49 | **0.001** |
| Age in years | -0.01 | 0.00 | -0.60 | 0.14 | -0.02 – -0.01 | -0.88 – -0.31 | -4.21 | **<0.001** |
| Sex [male] | -0.01 | 0.05 | -0.17 | 0.30 | -0.11 – 0.08 | -0.77 – 0.43 | -0.56 | 0.578 |
| Education in years | -0.01 | 0.01 | -0.13 | 0.14 | -0.03 – 0.01 | -0.43 – 0.16 | -0.94 | 0.355 |
| Observations | 46 | | | | | | | |
| $R^2$ / $R^2$ adjusted | 0.452 / 0.394 | | | | | | | |

Functional connectivity in older adults with plasma biomarkers was used as dependent variable, change in the AT-Term (1/ ($A\beta_{1-42}$/$A\beta_{1-40}$) * p-$tau_{217}$) from baseline to follow-up (on average 2 years and 8 months later), age in years at baseline, sex, and education in years were used as independent variables. AT-Term values were log-transformed. rsFC = resting-state functional connectivity. CI = 95% confidence interval.

### Table S12. Sample B: Linear model of effects of change in AT-Term on global efficiency

| | Network Global Efficiency | | | | | | | |
|---|---|---|---|---|---|---|---|---|
| *Predictors* | *Estimates* | *std. Error* | *std. Beta* | *standardized std. Error* | *CI* | *standardized CI* | *Statistic* | *p* |
| (Intercept) | 0.35 | 0.08 | 0.54 | 0.29 | 0.19 – 0.52 | -0.05 – 1.13 | 1.87 | **0.070** |
| Change in AT-Term | -0.02 | 0.01 | -1.49 | 0.36 | -0.04 – -0.01 | -2.22 – -0.75 | -4.09 | **<0.001** |
| Age in years | 0.00 | 0.00 | 0.25 | 0.15 | -0.00 – 0.00 | -0.06 – 0.56 | 1.66 | 0.106 |
| Sex [male] | 0.00 | 0.01 | 0.14 | 0.32 | -0.02 – 0.03 | -0.50 – 0.79 | 0.44 | 0.660 |
| Education in years | -0.00 | 0.00 | -0.16 | 0.15 | -0.01 – 0.00 | -0.47 – 0.15 | -1.05 | 0.300 |
| Observations | 46 | | | | | | | |
| $R^2$ / $R^2$ adjusted | 0.332 / 0.267 | | | | | | | |

The graph measure global efficiency in older adults with plasma biomarkers was used as dependent variable, change in the AT-Term (1/ ($A\beta_{1\text{-}42}$/$A\beta_{1\text{-}40}$) * p-$tau_{217}$) from baseline to follow-up (on average 2 years and 8 months later), age in years at baseline, sex, and education in years were used as independent variables. AT-Term values were log-transformed. CI = 95% confidence interval.

### Table S13. Sample C: Linear model of effects of temporal-lobe tau burden on functional connectivity

| | rsFC between left CA1 head - left subiculum head | | | | | | | | | |
|---|---|---|---|---|---|---|---|---|---|---|
| *Predictors* | *Estimates* | *std. Error* | *std. Beta* | *standardized std. Error* | *CI* | *standardized CI* | *Statistic* | *std. Statistic* | *p* | *std. p* |
| (Intercept) | 6.37 | 2.75 | 0.09 | 0.19 | 0.83 – 11.91 | -0.28 – 0.47 | 2.31 | 0.49 | **0.025** | 0.626 |
| Temporal-lobe tau PET burden | -3.22 | 1.41 | -0.13 | 0.13 | -6.05 – -0.39 | -0.40 – 0.14 | -2.29 | -0.94 | **0.027** | 0.351 |
| GFAP level | 0.09 | 0.04 | 0.58 | 0.15 | 0.01 – 0.16 | 0.28 – 0.87 | 2.31 | 3.94 | **0.025** | **<0.001** |

| | | | | | | | | | | |
|---|---|---|---|---|---|---|---|---|---|---|
| Age in years | -0.01 | 0.01 | -0.19 | 0.15 | -0.02 – 0.00 | -0.49 – 0.11 | -1.30 | -1.30 | 0.200 | 0.200 |
| Sex [male] | -0.04 | 0.06 | -0.20 | 0.27 | -0.16 – 0.07 | -0.73 – 0.34 | -0.74 | -0.74 | 0.465 | 0.465 |
| Education in years | 0.02 | 0.01 | 0.22 | 0.13 | -0.00 – 0.05 | -0.03 – 0.47 | 1.75 | 1.75 | 0.087 | 0.087 |
| Temporal-lobe tau PET burden × GFAP level | 0.01 | 0.00 | 0.60 | 0.15 | 0.00 – 0.01 | 0.31 – 0.89 | 4.13 | 4.13 | **<0.001** | **<0.001** |
| Observations | 55 | | | | | | | | | |
| $R^2$ / $R^2$ adjusted | 0.407 / 0.333 | | | | | | | | | |

Functional connectivity in older adults with tau PET was used as dependent variable, temporal-lobe tau PET burden, glial fibrillary acidic protein (GFAP) levels, age in years at baseline, sex, education in years, and the interaction of temporal-lobe tau PET burden by GFAP levels were used as independent variables. rsFC = resting-state functional connectivity. CI = 95% confidence interval. GFAP = glial fibrillary acidic protein.

### Table S14. Sample C: Linear model of effects of temporal-lobe tau burden on local efficiency

| | right ERC superficial local efficiency | | | | | | | |
|---|---|---|---|---|---|---|---|---|
| *Predictors* | *Estimates* | *std. Error* | *std. Beta* | *standardized std. Error* | *CI* | *standardized CI* | *Statistic* | *p* |
| (Intercept) | 2.82 | 0.77 | 0.15 | 0.23 | 1.27 – 4.37 | -0.31 – 0.62 | 3.68 | **0.001** |
| Temporal-lobe tau PET burden | -1.90 | 0.63 | -0.45 | 0.15 | -3.17 – -0.63 | -0.74 – -0.15 | -3.03 | **0.004** |
| Age in years | -0.01 | 0.01 | -0.20 | 0.15 | -0.02 – 0.00 | -0.51 – 0.10 | -1.37 | 0.177 |
| Sex [male] | -0.07 | 0.08 | -0.28 | 0.33 | -0.24 – 0.10 | -0.95 – 0.40 | -0.83 | 0.413 |
| Education in years | 0.00 | 0.02 | 0.02 | 0.15 | -0.03 – 0.04 | -0.29 – 0.33 | 0.16 | 0.873 |
| Observations | 45 | | | | | | | |
| $R^2$ / $R^2$ adjusted | 0.224 / 0.146 | | | | | | | |

The graph measure local efficiency in older adults with tau PET was used as dependent variable, temporal-lobe tau PET burden, age in years at baseline, sex, and education in years were used as independent variables. ERC = entorhinal cortex. CI = 95% confidence interval. *p*-FDR for temporal-lobe tau burden was 0.091.

### Table S15. Sample C: Linear model of effects of RSC tau burden on functional connectivity

| | rsFC between right tail (CA1) - left superficial RSC | | | | | | | |
|---|---|---|---|---|---|---|---|---|
| *Predictors* | *Estimates* | *std. Error* | *std. Beta* | *standardized std. Error* | *CI* | *standardized CI* | *Statistic* | *p* |
| (Intercept) | -1.63 | 0.52 | -0.36 | 0.20 | -2.67 – -0.58 | -0.76 – 0.04 | -3.12 | **0.003** |
| RSC tau PET burden | 1.67 | 0.39 | 0.58 | 0.14 | 0.88 – 2.47 | 0.31 – 0.85 | 4.26 | **<0.001** |
| Age in years | 0.00 | 0.00 | 0.12 | 0.13 | -0.00 – 0.01 | -0.13 – 0.37 | 0.93 | 0.356 |
| Sex [male] | 0.12 | 0.05 | 0.64 | 0.28 | 0.01 – 0.23 | 0.07 – 1.21 | 2.27 | **0.028** |
| Education in years | -0.00 | 0.01 | -0.02 | 0.14 | -0.03 – 0.02 | -0.30 – 0.25 | -0.17 | 0.866 |
| Observations | 55 | | | | | | | |
| $R^2$ / $R^2$ adjusted | 0.282 / 0.225 | | | | | | | |

Functional connectivity in older adults with tau PET was used as dependent variable, retrosplenial cortex (RSC) tau PET burden, age in years at baseline, sex, and education in years were used as independent variables. rsFC = resting-state functional connectivity. CI = 95% confidence interval. RSC = retrosplenial cortex.

### Table S16. Sample C: Linear model of effects of RSC tau burden on global efficiency

| | left superficial RSC global efficiency | | | | | | | |
|---|---|---|---|---|---|---|---|---|
| *Predictors* | *Estimates* | *std. Error* | *std. Beta* | *standardized std. Error* | *CI* | *standardized CI* | *Statistic* | *p* |
| (Intercept) | 0.14 | 0.15 | 0.19 | 0.19 | -0.17 – 0.45 | -0.19 – 0.57 | 0.91 | 0.368 |
| RSC tau PET burden | 0.44 | 0.12 | 0.50 | 0.13 | 0.21 – 0.68 | 0.24 – 0.75 | 3.84 | **<0.001** |
| Age in years | 0.00 | 0.00 | 0.09 | 0.12 | -0.00 – 0.00 | -0.15 – 0.33 | 0.73 | 0.467 |
| Sex [male] | -0.02 | 0.02 | -0.34 | 0.27 | -0.05 – 0.01 | -0.88 – 0.21 | -1.25 | 0.218 |
| Education in years | -0.00 | 0.00 | -0.06 | 0.13 | -0.01 – 0.01 | -0.32 – 0.20 | -0.47 | 0.637 |

| | |
|---|---|
| Observations | 55 |
| $R^2$ / $R^2$ adjusted | 0.355 / 0.303 |

The graph measure global efficiency in older adults with tau PET was used as dependent variable, retrosplenial cortex (RSC) tau PET burden, age in years at baseline, sex, and education in years were used as independent variables. rsFC = resting-state functional connectivity. CI = 95% confidence interval. RSC = retrosplenial cortex. *p*-FDR for RSC tau burden was 0.014.

### Table S17. Sample C: Linear model of effects of RSC tau burden on characteristic path length

| | **left superficial RSC characteristic path length** | | | | | | | |
|---|---|---|---|---|---|---|---|---|
| *Predictors* | *Estimates* | *std. Error* | *std. Beta* | *standardized std. Error* | *CI* | *standardized CI* | *Statistic* | *p* |
| (Intercept) | 4.55 | 0.71 | 0.05 | 0.19 | 3.12 – 5.97 | -0.33 – 0.43 | 6.40 | **<0.001** |
| RSC tau PET burden | -2.50 | 0.54 | -0.60 | 0.13 | -3.58 – -1.43 | -0.86 – -0.34 | -4.67 | **<0.001** |
| Age in years | -0.01 | 0.01 | -0.14 | 0.12 | -0.02 – 0.00 | -0.38 – 0.09 | -1.21 | 0.231 |
| Sex [male] | -0.02 | 0.07 | -0.09 | 0.27 | -0.17 – 0.12 | -0.63 – 0.45 | -0.33 | 0.742 |
| Education in years | 0.01 | 0.02 | 0.10 | 0.13 | -0.02 – 0.04 | -0.16 – 0.36 | 0.77 | 0.445 |
| Observations | 55 | | | | | | | |
| $R^2$ / $R^2$ adjusted | 0.361 / 0.309 | | | | | | | |

The graph measure characteristic path length in older adults with tau PET was used as dependent variable, retrosplenial cortex (RSC) tau PET burden, age in years at baseline, sex, and education in years were used as independent variables. rsFC = resting-state functional connectivity. CI = 95% confidence interval. RSC = retrosplenial cortex. *p*-FDR for RSC tau burden was < 0.001.

### Table S18. Sample A: Linear model of effects of functional connectivity on episodic memory

| | **Episodic memory performance** | | | | | | | |
|---|---|---|---|---|---|---|---|---|
| *Predictors* | *Estimates* | *std. Error* | *std. Beta* | *standardized std. Error* | *CI* | *standardized CI* | *Statistic* | *p* |
| (Intercept) | 0.30 | 0.13 | 0.35 | 0.20 | 0.05 – 0.56 | -0.06 – 0.75 | 2.41 | **0.020** |

| | | | | | | | | |
|---|---|---|---|---|---|---|---|---|
| rsFC left superficial CA3 body - left deep CA3 body | 0.20 | 0.07 | 0.32 | 0.12 | 0.05 – 0.35 | 0.09 – 0.56 | 2.75 | **0.008** |
| age | -0.29 | 0.07 | -0.47 | 0.12 | -0.44 – -0.14 | -0.70 – -0.23 | -3.95 | **<0.001** |
| *APOE4* Group [carrier] | -0.29 | 0.22 | -0.47 | 0.35 | -0.73 – 0.14 | -1.17 – 0.23 | -1.35 | 0.183 |
| sex [m] | -0.31 | 0.16 | -0.49 | 0.26 | -0.63 – 0.02 | -1.01 – 0.03 | -1.89 | 0.064 |
| education | 0.26 | 0.08 | 0.42 | 0.13 | 0.11 – 0.42 | 0.17 – 0.67 | 3.37 | **0.001** |
| Observations | 56 | | | | | | | |
| $R^2$ / $R^2$ adjusted | 0.370 / 0.307 | | | | | | | |

Episodic memory performance in amyloid and tau negative older adults was used as dependent variable, rsFC, age in years at baseline, *APOE4* group, sex, and education in years were used as independent variables. rsFC = resting-state functional connectivity. CI = 95% confidence interval.

### Table S19. Sample C: Linear model of effects of temporal-lobe tau burden on episodic memory

| | **Episodic memory performance** | | | | | | | |
|---|---|---|---|---|---|---|---|---|
| *Predictors* | *Estimates* | *std. Error* | *std. Beta* | *standardized std. Error* | *CI* | *standardized CI* | *Statistic* | *p* |
| (Intercept) | 0.20 | 0.14 | 0.30 | 0.21 | -0.08 – 0.47 | -0.11 – 0.72 | 1.44 | 0.157 |
| Temporal-lobe tau PET burden | -0.19 | 0.09 | -0.28 | 0.13 | -0.36 – -0.01 | -0.55 – -0.02 | -2.16 | **0.036** |
| Age in years | -0.14 | 0.09 | -0.22 | 0.13 | -0.31 – 0.03 | -0.48 – 0.05 | -1.65 | 0.106 |
| Sex [male] | -0.35 | 0.19 | -0.53 | 0.29 | -0.73 – 0.04 | -1.11 – 0.05 | -1.82 | 0.074 |
| Education in years | 0.24 | 0.09 | 0.37 | 0.14 | 0.06 – 0.42 | 0.09 – 0.64 | 2.63 | **0.011** |
| Observations | 54 | | | | | | | |
| $R^2$ / $R^2$ adjusted | 0.230 / 0.167 | | | | | | | |

Episodic memory performance in older adults with tau PET was used as dependent variable, temporal-lobe tau PET burden, age in years at baseline, sex, and education in years were used as independent variables. rsFC = resting-state functional connectivity. CI = 95% confidence interval.

### Table S20. Sample C: Linear model of interaction effect of functional connectivity by temporal-lobe tau burden on episodic memory

| | Episodic memory performance | | | | | | | |
|---|---|---|---|---|---|---|---|---|
| *Predictors* | *Estimates* | *std. Error* | *std. Beta* | *standardized std. Error* | *CI* | *standardized CI* | *Statistic* | *p* |
| (Intercept) | 0.30 | 0.13 | 0.41 | 0.21 | 0.04 – 0.57 | -0.01 – 0.82 | 2.29 | **0.026** |
| Temporal-lobe tau PET burden | -0.13 | 0.09 | -0.20 | 0.13 | -0.30 – 0.04 | -0.47 – 0.07 | -1.52 | 0.136 |
| rsFC left CA1 head - left subiculum head | -0.03 | 0.09 | -0.05 | 0.13 | -0.20 – 0.14 | -0.32 – 0.22 | -0.38 | 0.705 |
| Age in years | -0.18 | 0.09 | -0.28 | 0.14 | -0.36 – 0.00 | -0.57 – 0.00 | -1.99 | 0.052 |
| Sex [male] | -0.45 | 0.18 | -0.71 | 0.29 | -0.83 – -0.08 | -1.29 – -0.13 | -2.46 | **0.018** |
| Education in years | 0.28 | 0.09 | 0.44 | 0.14 | 0.10 – 0.46 | 0.15 – 0.72 | 3.06 | **0.004** |
| rsFC left CA1 head - left subiculum head × temporal-lobe tau PET burden | 0.23 | 0.08 | 0.35 | 0.12 | 0.07 – 0.38 | 0.11 – 0.60 | 2.88 | **0.006** |
| Observations | 54 | | | | | | | |
| $R^2$ / $R^2$ adjusted | 0.296 / 0.206 | | | | | | | |

Episodic memory performance in older adults with tau PET was used as dependent variable, temporal-lobe tau PET burden, rsFC, age in years at baseline, sex, education in years, and the interaction of temporal-lobe tau PET burden by rsFC were used as independent variables. rsFC = resting-state functional connectivity. CI = 95% confidence interval.

### Table S21. Sample C: Linear model of effects of temporal-lobe tau burden on episodic memory with low left CA1 head - left subiculum head rsFC

| | Episodic memory performance | | | | | | | |
|---|---|---|---|---|---|---|---|---|
| *Predictors* | *Estimates* | *std. Error* | *std. Beta* | *standardized std. Error* | *CI* | *standardized CI* | *Statistic* | *p* |
| (Intercept) | 0.31 | 0.22 | 0.39 | 0.31 | -0.14 – 0.75 | -0.25 – 1.03 | 1.42 | 0.169 |
| Temporal-lobe tau PET burden | -0.31 | 0.15 | -0.45 | 0.21 | -0.62 – -0.00 | -0.89 – -0.00 | -2.09 | **0.048** |

| | | | | | | | | |
|---|---|---|---|---|---|---|---|---|
| Age in years | -0.17 | 0.14 | -0.25 | 0.21 | -0.47 – 0.13 | -0.68 – 0.18 | -1.20 | 0.242 |
| Sex [male] | -0.43 | 0.28 | -0.62 | 0.40 | -1.01 – 0.15 | -1.45 – 0.22 | -1.53 | 0.139 |
| Education in years | 0.26 | 0.13 | 0.38 | 0.19 | -0.01 – 0.54 | -0.02 – 0.77 | 1.97 | 0.061 |
| Observations | 27 | | | | | | | |
| $R^2$ / $R^2$ adjusted | 0.288 / 0.158 | | | | | | | |

Episodic memory performance in older adults with tau PET and low rsFC (via a median-split) was used as dependent variable, temporal-lobe tau PET burden, age in years at baseline, sex, and education in years were used as independent variables. rsFC = resting-state functional connectivity. CI = 95% confidence interval.

### Table S22. Sample C: Linear model of effects of temporal-lobe tau burden on episodic memory with high left CA1 head - left subiculum head rsFC

| | Episodic memory performance | | | | | | | |
|---|---|---|---|---|---|---|---|---|
| *Predictors* | *Estimates* | *std. Error* | *std. Beta* | *standardized std. Error* | *CI* | *standardized CI* | *Statistic* | *p* |
| (Intercept) | 0.29 | 0.19 | 0.41 | 0.33 | -0.11 – 0.69 | -0.27 – 1.09 | 1.50 | 0.149 |
| Temporal-lobe tau PET burden | 0.08 | 0.12 | 0.13 | 0.20 | -0.17 – 0.33 | -0.29 – 0.55 | 0.65 | 0.520 |
| Age in years | -0.09 | 0.13 | -0.15 | 0.22 | -0.35 – 0.17 | -0.60 – 0.29 | -0.72 | 0.481 |
| Sex [male] | -0.47 | 0.31 | -0.79 | 0.52 | -1.10 – 0.17 | -1.87 – 0.29 | -1.52 | 0.144 |
| Education in years | 0.23 | 0.14 | 0.39 | 0.24 | -0.06 – 0.52 | -0.10 – 0.89 | 1.65 | 0.112 |
| Observations | 27 | | | | | | | |
| $R^2$ / $R^2$ adjusted | 0.202 / 0.057 | | | | | | | |

Episodic memory performance in older adults with tau PET and high rsFC (via a median-split) was used as dependent variable, temporal-lobe tau PET burden, age in years at baseline, sex, and education in years were used as independent variables. rsFC = resting-state functional connectivity. CI = 95% confidence interval.

### Table S23. Sample C: Linear model of interaction effect of functional connectivity on change in episodic memory

| | Change in episodic memory performance | | | | | | | |
|---|---|---|---|---|---|---|---|---|
| *Predictors* | *Estimates* | *std. Error* | *std. Beta* | *standardized std. Error* | *CI* | *standardized CI* | *Statistic* | *p* |
| (Intercept) | 0.11 | 0.11 | 0.17 | 0.22 | -0.12 – 0.34 | -0.28 – 0.62 | 0.94 | 0.351 |
| Temporal-lobe tau PET burden | -0.03 | 0.07 | -0.06 | 0.14 | -0.17 – 0.11 | -0.34 – 0.23 | -0.41 | 0.681 |
| rsFC left CA1 head - left subiculum head | -0.19 | 0.07 | -0.38 | 0.13 | -0.32 – -0.06 | -0.65 – -0.11 | -2.88 | **0.006** |
| *APOE4* Group [carrier] | 0.21 | 0.16 | 0.42 | 0.31 | -0.11 – 0.54 | -0.21 – 1.05 | 1.34 | 0.189 |
| Age in years | -0.02 | 0.07 | -0.03 | 0.14 | -0.15 – 0.12 | -0.32 – 0.25 | -0.23 | 0.818 |
| Sex [male] | -0.27 | 0.15 | -0.53 | 0.30 | -0.58 – 0.04 | -1.13 – 0.08 | -1.75 | 0.087 |
| Education in years | 0.25 | 0.07 | 0.50 | 0.15 | 0.11 – 0.40 | 0.21 – 0.80 | 3.46 | **0.001** |
| Observations | 45 | | | | | | | |
| $R^2$ / $R^2$ adjusted | 0.355 / 0.253 | | | | | | | |

Change in episodic memory performance in older adults with tau PET was used as dependent variable, temporal-lobe tau PET burden, rsFC, *APOE4* group, age in years at baseline, sex, and education in years were used as independent variables. rsFC = resting-state functional connectivity. CI = 95% confidence interval.

### Table S24. Sample C: Linear model of effect of Alzheimer's pathology on hippocampus volume

| | Bilateral TIV-corrected hippocampus volume | | | | | | | |
|---|---|---|---|---|---|---|---|---|
| *Predictors* | *Estimates* | *std. Error* | *std. Beta* | *standardized std. Error* | *CI* | *standardized CI* | *Statistic* | *p* |
| (Intercept) | 5812.81 | 123.33 | -0.05 | 0.26 | 5564.71 – 6060.91 | -0.58 – 0.47 | 47.13 | **<0.001** |
| Temporal-lobe tau PET burden | -12.33 | 67.31 | -0.03 | 0.14 | -147.74 – 123.08 | -0.32 – 0.26 | -0.18 | 0.855 |

| | | | | | | | | |
|---|---|---|---|---|---|---|---|---|
| AT-Term | -55.81 | 74.75 | -0.13 | 0.17 | -206.19 – 94.56 | -0.46 – 0.21 | -0.75 | 0.459 |
| GFAP | 87.64 | 94.27 | 0.18 | 0.19 | -102.01 – 277.29 | -0.21 – 0.57 | 0.93 | 0.357 |
| *APOE4* Group [carrier] | -18.69 | 180.73 | -0.04 | 0.39 | -382.26 – 344.89 | -0.82 – 0.74 | -0.10 | 0.918 |
| Age in years | -168.08 | 97.98 | -0.33 | 0.19 | -365.18 – 29.02 | -0.71 – 0.06 | -1.72 | 0.093 |
| Sex [male] | 49.21 | 157.17 | 0.11 | 0.34 | -266.96 – 365.39 | -0.57 – 0.78 | 0.31 | 0.756 |
| Education in years | -105.17 | 76.65 | -0.22 | 0.16 | -259.36 – 49.02 | -0.53 – 0.10 | -1.37 | 0.177 |
| Observations | 55 | | | | | | | |
| $R^2$ / $R^2$ adjusted | 0.126 / -0.005 | | | | | | | |

Bilateral total intracranial volume (TIV)--corrected hippocampus volume in older adults with tau PET was used as dependent variable, temporal-lobe tau PET burden, AT-Term, GFAP level, *APOE4* group, age in years at baseline, sex, and education in years were used as independent variables. CI = 95% confidence interval.

### S25. Within-cohort replication of associations discovered in resting-state data using breathing-task data

| Model | Functional connection/ graph measure | beta | 95% CI | df | t | *p* |
|---|---|---|---|---|---|---|
| **Age** | left BA36 - left DG body | -0.06 | -0.35, 0.22 | 50 | -0.45 | 0.66 |
| | right deep subiculum body - left BA36 | -0.09 | -0.37, 0.19 | 50 | -0.67 | 0.51 |
| | network local efficiency | -0.21 | -0.95, 0.24 | 50 | -0.86 | 0.30 |
| | nodal local efficiency of left CA3 head | -0.24 | -0.57, 0.09 | 43 | -1.48 | 0.15 |
| **Age * *APOE4* Group** | left CA3 superficial body - left CA3 deep body | 0.69 | -0.17, 1.54 | 49 | 1.62 | 0.11 |
| **AT-Term** | right BA36 - left CA1 head | 0.13 | -0.12, 0.39 | 63 | 1.06 | 0.29 |

| | | | | | | |
|---|---|---|---|---|---|---|
| **Change in AT-Term** | network global efficiency | -0.66 | -0.94, -0.38 | 39 | -4.71 | < 0.001 |
| | nodal global efficiency left tail (CA1) | -0.17 | -0.50, 0.16 | 39 | -1.05 | 0.298 |
| | nodal global efficiency left DG body | -0.19 | -0.56, 0.17 | 39 | -1.07 | 0.290 |
| | nodal global efficiency left CA1 deep body | -0.23 | -0.59, 0.13 | 39 | -1.28 | 0.206 |
| | nodal global efficiency left deep BA35 | -0.61 | -0.91, -0.30 | 39 | -4.00 | < 0.001 |
| | nodal characteristic path length right superficial BA35 | 0.16 | -0.21, 0.52 | 39 | 0.88 | 0.386 |
| **Temporal-lobe tau * GFAP** | left CA1 head - left subiculum head | 0.16 | -0.20, 0.53 | 45 | 0.91 | 0.368 |
| | left BA36 - left CA1 head | 0.31 | -0.03, 0.65 | 45 | 1.81 | 0.077 |
| | left superficial PHC - left superficial ERC | 0.18 | -0.16, 0.53 | 45 | 0.88 | 0.431 |
| | left tail (CA1) - left subiculum head | 0.37 | 0.02, 0.72 | 45 | 2.11 | 0.040 |
| | left BA36 - left superficial ERC | 0.30 | -0.06, 0.65 | 45 | 1.69 | 0.098 |
| **RSC tau** | | | | | | |
| | right tail (CA1) - left superficial RSC | 0.29 | -0.03, 0.60 | 47 | 1.84 | 0.072 |
| | left deep subiculum body - left deep RSC | 0.34 | 0.03, 0.64 | 47 | 2.18 | 0.034 |
| | right tail (CA1) - left deep RSC | 0.18 | -0.14, 0.50 | 47 | 1.14 | 0.262 |
| | nodal global efficiency of left superficial RSC | 0.19 | -0.13, 0.50 | 47 | 1.19 | 0.238 |

| | | | | | | |
|---|---|---|---|---|---|---|
| | nodal characteristic path length of left superficial RSC | -0.06 | -0.73, 0.62 | 47 | -0.17 | 0.869 |

For all functional connections that were part of significant clusters using the network-based statistics (NBS) approach and significant FDR-corrected graph measures in the resting-state data, the models were repeated in the breathing task data. Standardized betas, degrees of freedom (df), t-values (t), the corresponding 95% confidence interval (CI), and uncorrected p-values are reported. DG = dentate gyrus. PHC = parahippocampal cortex. ERC = entorhinal cortex. RSC = retrosplenial cortex.

### S26. Associations discovered in breathing-task data using cluster correction

| **Model** | **Functional connection/ graph measure** | **beta** | **95% CI** | **df** | **t** | ***p*-FDR** |
|---|---|---|---|---|---|---|
| **Age** | - | | | | | |
| **Age * *APOE4* Group** | - | | | | | |
| **AT-Term** | - | | | | | |
| **Change in AT-Term** Cluster *p*-FDR = 0.024 [95%$CI_p$ 0.021, 0.027] | left CA1 head - right superficial subiculum body | 0.58 | 0.29, 0.87 | 39 | 4.03 | - |
| Cluster *p*-FDR = 0.024 [95%$CI_p$ 0.021, 0.027] | right DG head - left CA1 head | 0.56 | 0.25, 0.87 | 39 | 3.62 | - |
| Cluster *p*-FDR = 0.024 [95%$CI_p$ 0.021, 0.027] | right DG head - right DG body | 0.53 | 0.21, 0.84 | 39 | 3.36 | - |
| | network global efficiency | -0.66 | -0.94, -0.38 | 39 | -4.71 | < 0.001 |
| | nodal global efficiency left BA36 | -0.63 | -0.91, -0.35 | 39 | -4.55 | 0.002 |
| | nodal global efficiency left deep BA35 | -0.61 | -0.91, -0.30 | 39 | -4.00 | 0.006 |
| | nodal global efficiency left superficial BA35 | -0.55 | -0.85, -0.25 | 39 | -3.72 | 0.008 |

| | | | | | | |
|---|---|---|---|---|---|---|
| | nodal global efficiency right superficial RSC | -0.57 | -0.88, -0.25 | 39 | -3.66 | 0.008 |
| | nodal global efficiency left superficial ERC | -0.50 | -0.80, -0.19 | 39 | -3.29 | 0.018 |
| | nodal global efficiency left superficial CA3 body | -0.51 | -0.83 – -0.19 | 39 | -3.22 | 0.018 |
| | nodal global efficiency left superficial RSC | -0.50 | -0.83 – -0.17 | 39 | -3.09 | 0.022 |
| | nodal global efficiency left deep RSC | -0.50 | -0.83 – -0.17 | 39 | -3.05 | 0.022 |
| | nodal global efficiency right deep CA3 body | -0.44 | -0.77 – -0.12 | 39 | -2.76 | 0.041 |
| | nodal global efficiency right deep RSC | -0.44 | -0.77 – -0.11 | 39 | -2.69 | 0.044 |
| | nodal global efficiency right deep BA35 | -0.44 | -0.77 – -0.10 | 39 | -2.65 | 0.044 |
| **Temporal-lobe tau * GFAP** Cluster $p$-FDR = 0.022 [95%CI$_p$ 0.019, 0.025] | left tail (CA1) - left deep subiculum body | 0.61 | 0.30, 0.91 | 45 | 3.95 | - |
| Cluster $p$-FDR = 0.022 [95%CI$_p$ 0.019, 0.025] | left tail (CA1) - left superficial subiculum body | 0.62 | 0.30, 0.93 | 45 | 3.90 | - |
| **RSC tau** | - | | | | | |

For all functional connections for the breathing-task data that were part of significant clusters using the network-based statistics (NBS) approach, FDR-corrected cluster p-value and the corresponding 95% confidence interval (CI), standardized betas, degrees of freedom (df), t-values (t), and the corresponding 95% confidence interval (CI) are reported. The same applies to significant graph measures, with FDR-corrected p-values per region (network-level measures are tested once and therefore FDR-correction is not applied there). DG = dentate gyrus. RSC = retrosplenial cortex. ERC = entorhinal cortex.

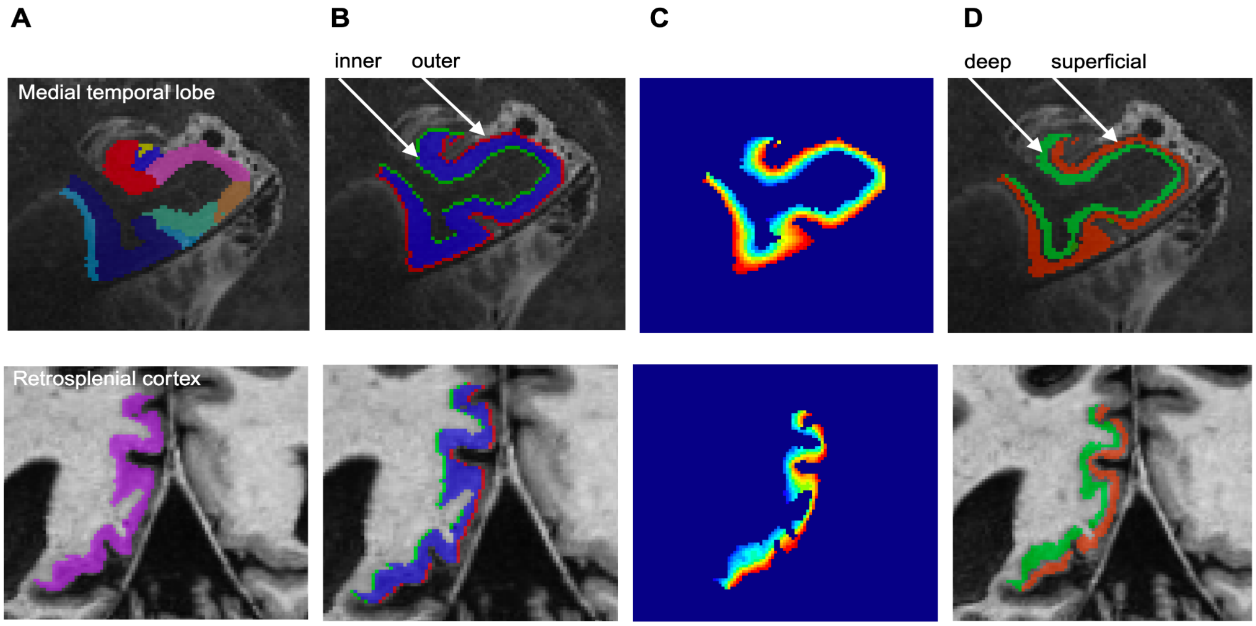


**Figure S1. Layer processing pipeline. A)**. Upper row: ASHS segmentation of the 7 Tesla T2-weighted image was used to derive medial temporal lobe (MTL) ROIs. Lower row: Fastsurfer DKT atlas segmentation of the 7 Tesla T1-weighted image was used to derive a retrosplenial cortex (RSC) isthmus cingulate ROI. **B)** For the whole MTL without dentage gyrus and for RSC, a geometrical approach was used to define the inner (gray to white matter; green) and outer (gray matter to CSF; red) border per slice. **C)** LayNii was used to derive 20 equidistant layers based on prior research. **D)** Layers 2 - 9 were binned to a deep layer (light green segmentation), layers 12 - 19 were binned to a superficial layer (light red segmentation). Not pictured: This was then overlaid with the separate ASHS MTL ROI masks. Layers were derived in the hippocampal body for CA1, CA3, subiculum, as well as for BA35, entorhinal cortex, parahippocampal cortex, and RSC, where layers passed visual inspection. Deep layers of CA1, CA3, and subiculum refer to the "inner" layer segment likely covering pyramidal layers and superficial layers refer to the "outer" segment likely covering stratum lacunosum moleculare.

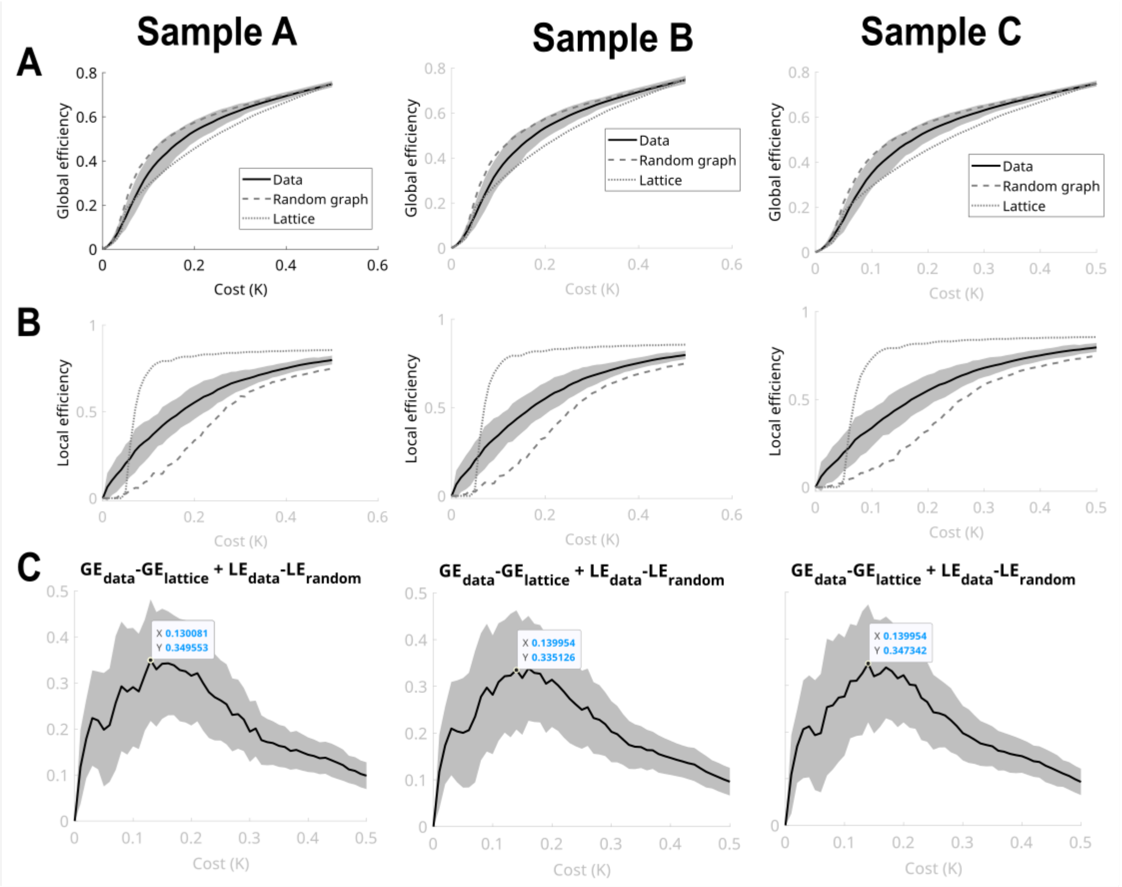


**Figure S2. Optimal cost threshold determination for sample A, B, and C. A)**. Global efficiency (GE) is plotted across varying cost thresholds, comparing how the real network (data) behaves relative to random and lattice structures. The random network acts as a baseline for randomization effects, while the lattice represents a highly ordered structure. **B)** Local efficiency (LE) is plotted across cost thresholds for the real, random, and lattice network. **C)** The optimal cost threshold is determined by maximizing the difference between global and local efficiency values for real data compared to lattice and random networks, reflecting the network's small-world configuration. We used the formula *GE(Data) - GE(Lattice) + LE(Data) - LE(Random)*. The highest value on the y-axis is used as cost threshold, as it indicates the maximum divergence between the real and the modeled networks, with the real network's distinctive properties being most prominent. This resulted in an optimal cost threshold of 0.13 for sample A, 0.14 for sample B, and 0.14 for sample C. The cost threshold of 0.14, for example, means that the 14% largest correlation coefficient values were used to build the binary adjacency matrix, which either uses 0 or 1 to define whether there is a suprathreshold connection between the respective pair of nodes. See (Achard & Bullmore, 2007) for details.

### References Supplementary:


Achard, S., & Bullmore, E. (2007). Efficiency and cost of economical brain functional networks. *PLoS Computational Biology*, *3*(2), e17.
